# Sharp habitat shifts, evolutionary tipping points and rescue: quantifying the perilous path of a specialist species toward a refugium in a changing environment

**DOI:** 10.1101/2023.09.28.559956

**Authors:** Léonard Dekens

## Abstract

Specialists species thriving under specific environmental conditions in narrow geographic ranges are widely recognised as heavily threatened by climate deregulation. Many might rely on both their potential to adapt and to disperse toward a refugium to avoid extinction. It is thus crucial to understand the influence of environmental conditions on the unfolding process of adaptation. Here, I study the eco-evolutionary dynamics of a sexually reproducing specialist species in a two-patch quantitative genetic model with moving optima. Thanks to a separation of ecological and evolutionary time scales and the phase-line study of the selection gradient, I derive the critical environmental speed for persistence, which reflects how the existence of a refugium impacts extinction patterns. Moreover, the analysis provides key insights about the dynamics that arise on the path towards this refugium. I show that after an initial increase of population size, there exists a critical environmental speed above which the species crosses a tipping point, resulting into an abrupt habitat switch. Besides, when selection for local adaptation is strong, this habitat switch passes through an evolutionary “death valley”, leading to a phenomenon related to evolutionary rescue, that can promote extinction for lower environmental speeds than the critical one.

## 1 Introduction

### Biological context

Historical data highlight how climate change shifts the spatial distributions of species across taxa, especially polewards (Parmesan et al. 1999 on butterflies) or upwards (Lenoir et al. 2008 on plants, Moritz et al. 2008 on mammals). Predicting the interplay between this change in species distribution and species abundance and persistence is an ongoing crucial challenge (Ehrlén and Morris 2015) that requires to better understand the evolutionary strategies of adaptation in face of climate change (Hoffmann and Sgrò 2011). One category of species that has been found to be particularly vulnerable to the changing climate gathers the various types of specialist species (Clavel, Julliard, and Devictor 2011). As these are adapted to a limited niche width, opportunities to disperse and adapt to a potentially more suitable habitat are sparser, especially in increasingly fragmented environments (Berg et al. 2010, Adams-Hosking et al. 2012, Hof, Jansson, and Nilsson 2012, Damschen et al. 2012). This issue highlights the importance of habitats that can act as refugia, which have already been shown to have played a major part in specialists’ persistence in the past (Corlett and Tomlinson 2020) and is recognised as a key component in conservation efforts (Morelli et al. 2016). These refugia can be thermal shelters located polewards to escape the rising temperatures, hydrological refugia in island-continent systems (McLaughlin et al. 2017, Ramirez et al. 2020) or edaphic refugia for soil specialists (Corlett and Tomlinson 2020). These examples emphasize the relevance of incorporating a spatial structure of patchiness in models of adaptation to climate change to better understand the population dynamics of specialists species towards refugia in highly fragmented environments (Urban et al. 2016).

### Theoretical models of adaptation to a moving optimum in a single habitat: lag-load

To understand and predict how species adapt to a changing environment, one can turn to theoretical models. The case of gradually moving environments has been attracting sustained interest from quantitative genetic models for over thirty years (see a review in [Kopp and Matuszewski 2014]). One of the first lines of research focused on the demographic and trait dynamics of a panmictic population living in a single habitat, subjected to stabilizing selection around an optimal trait moving at a constant speed (Lynch and Lande 1993; Bürger and Lynch 1995; Lande and Shannon 1996). The analysis highlights how maladaptation to the changing environment induces a lag between the population’ mean trait and the optimal trait, which eventually stabilizes. This evolutionary load impacts the demography by decreasing the population size. Therefore, in this case, these studies derive a simple expression of the critical rate of environmental change above which the environment changes too fast for the population to persist and leads to extinction. Key features underlying the analytical results of these studies resides in the following assumptions: a Gaussian approximation of the trait distribution within the population and a quadratic selection function implying that maladaptation increases more steeply away from the optimal trait. This approach has next been extended to include the effects of plasticity (Chevin, Lande, and Mace 2010), multidimensional quantitative traits (Duputié et al. 2012), age-structure (Cotto and Ronce 2014; Cotto, Sandell, et al. 2019), or to examine the influence of the mode of reproduction (Waxman and Peck 1999, Garnier et al. 2022).

### Evolutionary tipping points

However, a more recent study [Osmond and Klausmeier 2017] showed that this feature of a constant lag at equilibrium between the mean trait and the optimal trait is a consequence of the particular quadratic shape of the stabilizing selection function chosen in the references above, where the selection gradient increases linearly with the distance between the population’s mean trait and the moving optimum. For other selection functions where the strength of selection instead fades away from the moving optimum, [Osmond and Klausmeier 2017] showed that there exists evolutionary tipping points that, when crossed by increasing the rate of environmental change, abruptly lead the population to extinction. This happens because, for these selection functions leading to a change of convexity in the selection gradient, the lag grows indefinitely past a certain threshold. This feature (among many others) was also characterized analytically in [Garnier et al. 2022] that investigated more broadly the influence of the selection functions on the adaptation of sexually and asexually reproduction populations to a changing environment. However, while these evolutionary tipping points are linked to particular choices of selection functions in a single habitat framework, they have been reported to arise in more complex frameworks (Klausmeier et al. 2020). For example, including an age structure to the population (Cotto and Ronce 2014, Cotto, Sandell, et al. 2019) allows for feedback loops between the dynamics of the demography and the trait dynamics to create multiple co-existing equilibria that promote evolutionary tipping points. This last feature has also been related to tipping points in a broader variety of eco-evolutionary models (see Dakos et al. 2019): for example, abrupt switching between different developmental strategies in an oscillatory environment (Botero et al. 2015) or between ecosystem structure in shallow lake environments (Chaparro-Pedraza 2021).

### Spatial structure with changing environment

As spatial structure provides species with the possibility to disperse when facing a changing environment, several theoretical quantitative genetic studies have included a spatial component in their modelling. A particularly prolific line of research considers a species evolving in a continuous space, extending the concept of a gradually moving optimum in a single habitat to an environmental gradient shifting gradually at a constant speed. Stemming from the framework introduced by [Pease, Lande, and Bull 1989], more and more sophisticated models have analyzed how populations can track the shifting environmental gradient with a constant spatial lag when the speed of the environment is below a critical threshold, thus escaping extinction by shifting their spatial range. They have built extensions to study the influence density-dependence (Polechová, Barton, and Marion 2009), or a multidimensional adaptive trait (Duputié et al. 2012). More recently, a study modelling two dispersal modes differing in their mean dispersal distance (pollen and seed dispersal) shows that long range-dispersal can trigger an ecological niche shift in addition to the spatial range shift, which buffers the species for larger environmental speeds (Aguilée et al. 2016). All these analyses rely heavily on the analytical travelling waves toolkit that is specifically designed to study the long-term effect of dispersal in a continuous space (see Alfaro, Berestycki, and Raoul 2017, Roques et al. 2020 and Lavigne 2023 for precise mathematical expansion in the case of asexual populations).

However, these methods are not suited to study the patterns of dispersal in fragmented and patchy environments, where the demographic dynamics and the trait dynamics are quite difficult to disentangle, even under stable environment (see Ronce and Kirkpatrick 2001; Hendry, Day, and Taylor 2001; Dekens 2022 for sexual reproduction and Débarre, Ronce, and Gandon 2013; Mirrahimi 2017; Mirrahimi and Gandon 2020 for asexual ones). Therefore, most models studying adaptation to a changing and fragmented environment rely mostly on numerical simulations to explore complex metacommunities dynamics (see Cotto, Wessely, et al. 2017 for such a model with multiple traits, species and an age structure, and Walters and Berger 2019 on the contribution of genetic variance on the time to extinction in a migration-mutation-selection-drift framework), or to assess the interplay between dispersal and local competition under a warming climate (Thompson and Fronhofer 2019,McManus et al. 2021). Moreover, simulations of a quantitative genetic two-patch model suggest that a changing environment can first lead to sharp declines in subpopulation size with a potential rebound when it stabilizes, but less so for specialists (Bourne et al. 2014). As it is important to quantify these sharp dynamics and predict the conditions under which they occur, I propose here to analyze a two-patch quantitative genetic model under changing environment. It aims at improving our understanding of the evolutionary mechanisms of a specialist species, particularly how they can potentially leverage the existence of a refugium when their native habitat becomes non viable.

Indeed, under a changing environment, such a species, while initially being well-adapted to its native habitat, is expected to lag behind the optimum of the native habitat, and thus closer to the refugium’s one. Under which conditions do species keep pace with their native habitat? If the environmental speed is too strong, will they be strained to adapt to the refugium, and if so, how successfully? To answer these questions, I will, as a starting point, leverage the results of an analogous model under stable environment [Dekens 2022] that includes the analytical derivation of the source-sink dynamics characteristic of a specialist species that I use here as the initial state of the system.

### Outline of the paper

In this work, I study the eco-evo dynamics of a sexual population in a fragmented and changing environment thanks to a two-patch quantitative genetic model with moving optima. Precisely, I consider a specialist population initially adapted to one of the two habitats (their native habitat), whose migrants fail to establish in the other one at first (the refugium). My aim is to predict analytically the dynamics of niche evolution of this specialist population as a function of the speed of environmental change. To this aim, in Section 2, I show that, in a regime of small within-family variance allowing to separate ecological and evolutionary time scales, all the features of the dynamics of adaptation can be deduced from a simple modification of the phase-line analysis of the selection gradient derived in a previous study of a two-patch model under stable environment (Dekens 2022). Th results are presented in Section 3. First, for small environmental speeds, the metapopulation actually increases in size, as a result of a relative beneficial loss of specialization. Next, there exists an intermediate critical environmental speed leading to an abrupt habitat switch from the native habitat to the refugium. Such a switch corresponds to an evolutionary tipping point and is therefore difficult to reverse. Moreover, above a selection threshold leading to the creation of a evolutionary “death valley” between the habitats, the population experiences evolutionary rescue during the habitat switch. Finally, I quantify the critical speed of environmental change above which the population becomes too maladapted and goes extinct in this native habitat/refugium framework. Contrary to the classical prediction in a single-habitat framework, I show that, here, the critical speed is not always increasing with selection strength and can be discontinuous.

## 2 Methods

### 2.1 Model

The model follows a classical setting of quantitative genetic models for heterogeneous environments (Ronce and Kirkpatrick 2001; Hendry, Day, and Taylor 2001; Débarre, Ronce, and Gandon 2013; Mirrahimi and Gandon 2020; Dekens 2022), which is summarized in Fig. 1. It considers a sexually reproducing species that lives in a fragmented two-patch environment, where one patch represents the species’ native habitat to which it is specialized and the other a refugium to which the species is not initially adapted. These habitats are connected by a back-and-forth migration at a constant rate ***m*** *>* 0. Within each habitat, individuals mate randomly at a rate ***r*** and die from density-dependent regulation at a rate ***κ*** *>* 0 and trait-dependent stabilizing selection of intensity ***g*** *>* 0. The latter is minimal at a local trait optimum (***θ***_**2**_(***t***) in the native habitat and ***θ***_**1**_(***t***) in the refugium, with ***θ***_**1**_(***t***) *<* ***θ***_**2**_(***t***)) and decreases quadratically away from it. The two local optima are assumed to shift at the same constant speed ***c*** *>* 0, which models the action of the shifting environment: ***θ***_***i***_(***t***) = (−1)^*i*^***θ***+***ct***. Individuals are characterized by a quantitative trait *z* ∈ R that determines their adaptation to the habitat they live in. This quantitative trait is thought to have a highly polygenic basis with small additive allelic effects, and its inheritance across generations is modelled by the infinitesimal model (Fisher 1919; Bulmer 1971; Turelli and Barton 1990; Barton, Etheridge, and Véber 2017. The infinitesimal model in its additive version states that the distribution of trait within each family is Gaussian, centered on the mean parental trait and with variance ***σ***^2^, which reads

**Figure 1.**
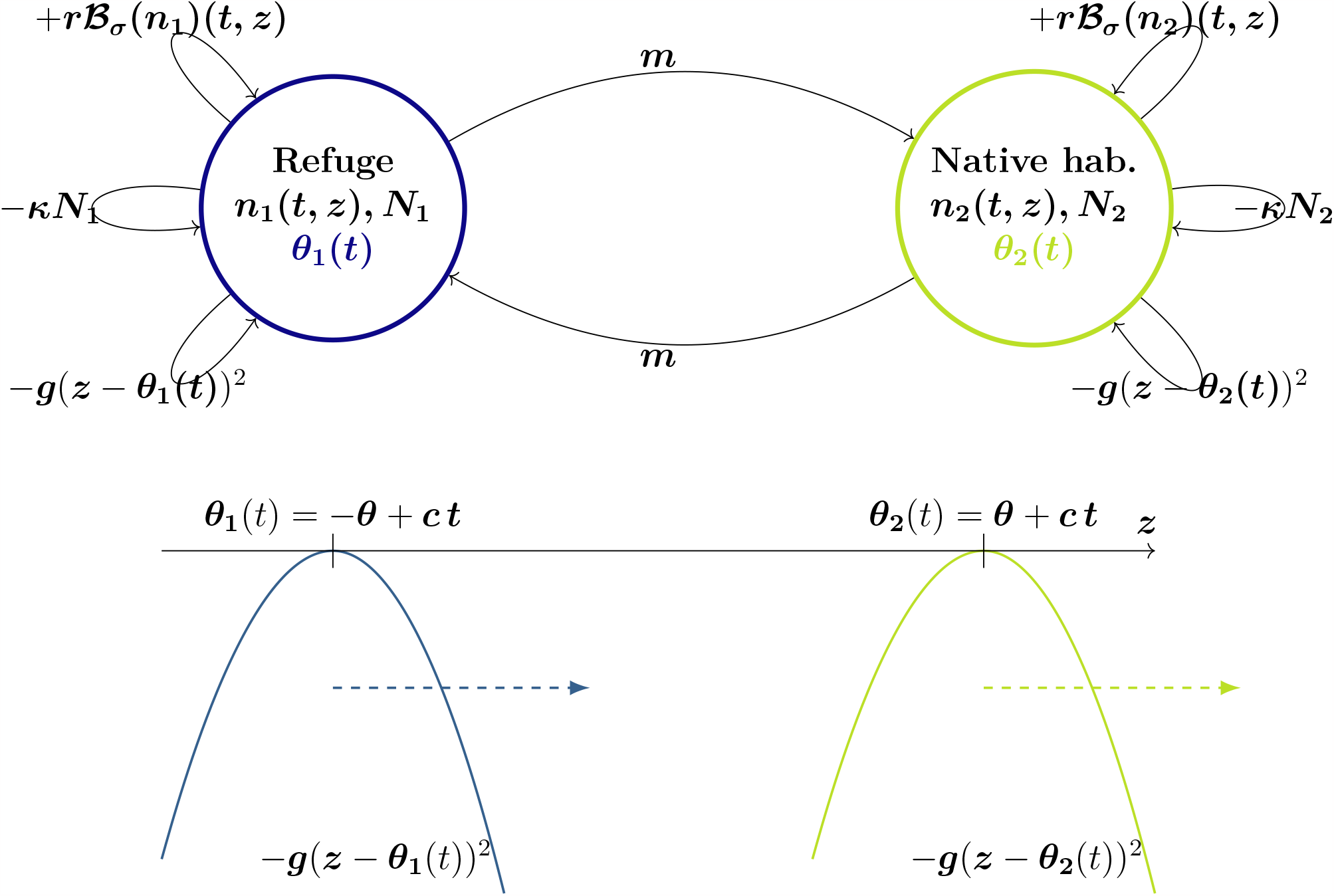
Two-patch changing environment framework for a quantitative trait*z*.

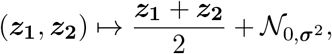

where ***z***_**1**_ and ***z***_**2**_ represent the two parental traits. The parameter ***σ***^2^ is the within-family variance (also called segregational variance). In this model, it is assumed to be constant across families, time and space. Accordingly, at a time ***t***, the number of individuals born with a trait ***z*** in the habitat *i* is given by the following formula (also used Turelli and Barton 1990; Mirrahimi and Raoul 2013; Calvez, Garnier, and Patout 2019; Patout 2020; Dekens and Lavigne 2021; Dekens 2022; Garnier et al. 2022):

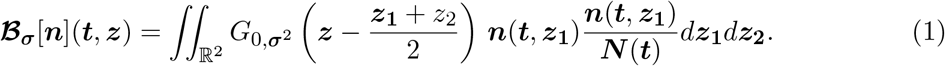

We consider the dynamics of the local trait distributions given by:

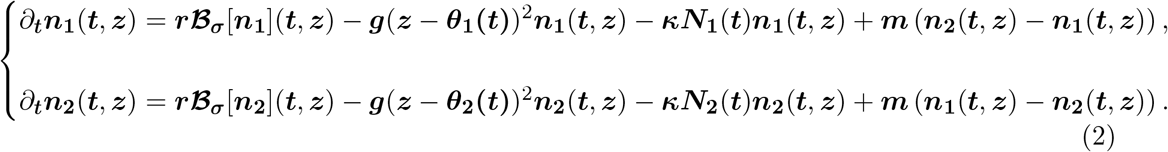

The environment changes through time at a constant speed ***c*** which is modelled by a linear increase of the local optima: ***θ***_**1**_ = −***θ*** + ***ct*** and ***θ***_**2**_ = ***θ*** + ***ct***.

### 2.2 Overview of the analysis

In this section, I explain how to conveniently transform the PDE system (2) in order to isolate the influence of the changing environment, which allows to leverage the main ideas of the analysis under stable environment done in [Dekens 2022]. In fact, I show that, in a chosen regime of small within-family variance, the full dynamics of the local trait distributions can be summarized by the dynamics of two time-dependent variables: the ratio of the two subpopulations sizes and the metapopulation’s mean trait. Moreover, I show that, in the final system *S*, the environmental change directly influences only the dynamics of the latter, and only linearly. I refer the interested reader who wishes to learn about all the mathematical details underlying this section to Appendix A.

1. **Small within-variance regime and moving-frame reference**. I place the analysis in the regime where the within-family variance ***σ***^2^ is small compared to the difference between the local optima (***θ***_**2**_−***θ***_**1**_ = 2***θ***). This is likely to be the case after a long time at equilibrium under stabilizing selection and stable environment. In this regime, the local genetic variances are expected to remain small, so local mean traits take a long time to shift under the action of local selection (see Dekens 2022). Therefore, it is practical to rescale the time to match the timescale of the evolution of the local mean traits. Moreover, it is also convenient to place the analysis in the moving-frame reference whose speed matches the environmental speed, in which the local optimal traits are fixed. In the first paragraph of Appendix A, I detail how to do so along with rescaling other variables and parameters to get a dimensionless system from (2). From now on, I will refer to these rescaled quantities, which will be not bolded.
2. **Initial specialized population**. In all what follows, I examine the case where selection is strong enough relative to migration for specialization to exist at equilibrium and be stable under a stable environment: 1 + 2*m <* 5*g* (see Proposition 4.2 of Dekens 2022). Moreover, I also assume that selection is upper bounded when migration is strong to ensure that this specialist equilibrium is viable, which reads: *g*(*m*−1) *< m*^2^ (see also Proposition 4.2 of Dekens 2022). Under these conditions, there exist two viable specialist equilibria according to mirrored source-sink dynamics. For the initial state of the system, I choose the equilibrium describing a species specialized in the native habitat whose precise characterization is indicated in the Proposition 4.2 of [Dekens 2022].
3. **Gaussian approximation of local trait distributions**. In the chosen regime of small within-family variance, arguments developed in [Dekens 2022] ensure that the local trait distributions are approximately Gaussian, with a small variance (twice the within-family variance). Therefore, the moment-based ODE system describing the dynamics of the subpopulations sizes and the local mean traits is closed (see (8)). I choose to report the analysis on this moment-based ODE system instead of the full PDE system on the local trait distributions (see the second paragraph of Appendix A for details about the derivation of such a moment-based system).
4. **Separation of time scales and limit system**. To disentangle the coupled dynamics of the subpopulations sizes and the local mean traits, one can leverage the fact that the local selection terms driving the dynamics of the mean traits are proportional to the small local genetic variances (see Eq. (8)). Therefore, there exists a separation of time scale between fast ecological phenomena (birth/death/migration) and slow evolutionary ones (shift of the local mean traits by selection), similarly as in the analysis of [Dekens 2022]. In the third paragraph of Appendix A, I show that this leads to a final system whose complexity is greatly reduced, as it only involves the ratio between subpopulations sizes 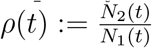 and the metapopulation’s mean trait *Z*(*t*). It reads

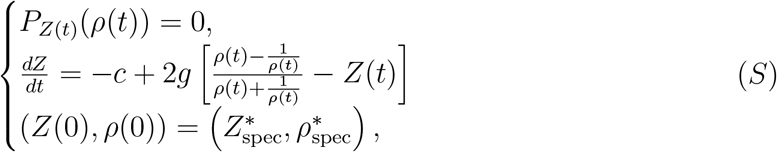

where *P*_*Z*_ is a third-order polynomial whose coefficients depend on *Z* (details can be found in Lemma 1 of Dekens 2022) and 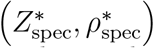 are the initial values defined in A.

The keen reader might notice that *S* is almost the same as the analogous one derived in [Dekens 2022] under stable environment. The only but crucial difference is that here, the changing environment pushes the metapopulation’s mean trait *Z*(*t*) backwards with a speed −*c*, which results in a lag (remember that the analysis is done in the moving-frame reference). Although this could look like a rather benign change, it leads to very rich dynamics that I detail in Section 3. In the next subsection, I explain how to use *S* in a simple way to predict these dynamics.

### 2.3 Predicting the new equilibrium under a changing environment with a phase lines’ study

Consider the first line of *S*. For a given average trait in the metapopulation *Z*(*t*), it states that *ρ*(*t*) is a (positive) root of *P*_*Z*(*t*)_. In the case where such a root is uniquely defined, *ρ*(*t*) can be seen as a well-defined function of *Z*(*t*) which I thus note *ρ*(*Z*(*t*)) (see Section 3 of [Dekens 2022] for the analytical conditions under which this happens). In this case, one can circumvent the first algebraic equation of *S* to reduce the analysis to to the following differential equation only on *Z*(*t*):

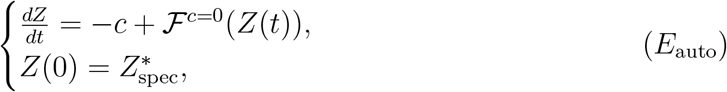

where I define the function 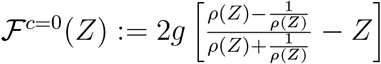 that does not depend directly on the environmental change. Biologically, the function ℱ^*c*=0^ can be interpreted as the **selection gradient under stable environment** (*c* = 0; note that the trait variance does not appear because it it scaled to 1 in the time scale considered). It represents the selection force pushing the mean trait *Z*(*t*) towards an optimum which integrates the demographic feedback on the ecological dynamics on the evolutionary ones. Precisely, this optimum depends on the balance between the two subpopulation sizes 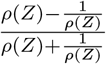 (I recall that *ρ*(*Z*) is the ratio between the subpopulation sizes when the mean trait is *Z*).

The differential equation *E*_auto_ does not involve time directly as a variable, so it is qualified as autonomous. The dynamics described by such an equation are conveniently studied through its phase line, which is the graph of the function *Z↦ − c* + ℱ^*c*=0^(*Z*) (right-hand side of (*E*_auto_)). When it is positive (resp. negative), *Z*(*t*) increases (resp. decreases), so the equilibria are located where it cancels (they are stable when the slope is negative and unstable when it is positive). Notice that the only impact of the changing environment is only a vertical translation of the phase-line under stable environment (positive values of *c* correspond to downward translations). This implies that it can drive some stable-environment equilibria to disappear or to shift. More precisely, the new equilibrium obtained from the initial specialist state is located at the rightmost intersection of the downward-shifted phase line and the *x*-axis that has a negative slope (see Fig. 2 for an illustration).

**Figure 2.**
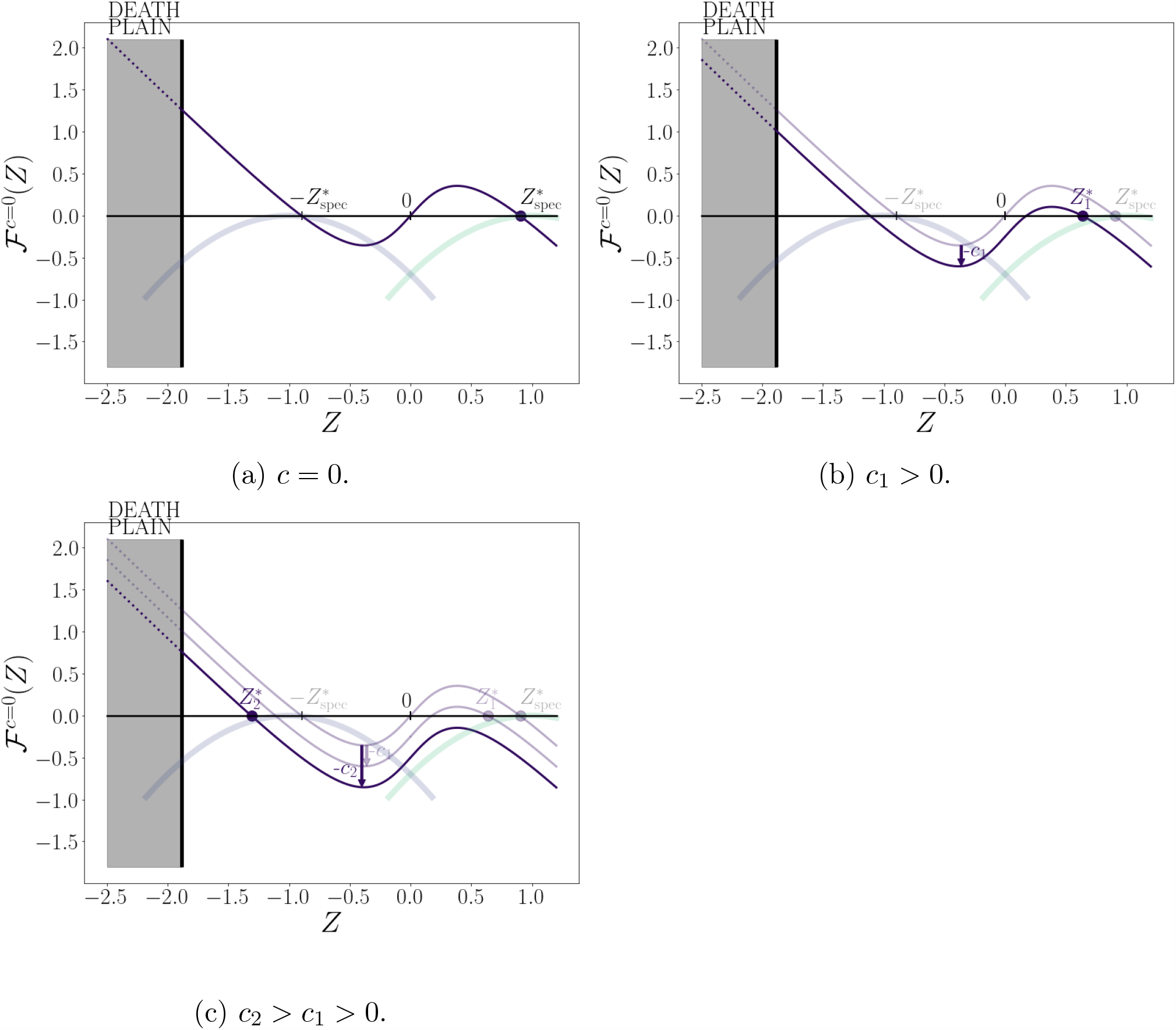
Illustration of the selection gradient’s phase line (purple dark line) (*E*_auto_) for several environmental speeds. (top left: stable environment *c* = 0, top right: environmental speed *c*_1_ *>* 0, bottom left: environmental speed *c*_2_ *> c*_1_ *>* 0; for all subfigures: *g* = 0.7, *m* = 0.5). The local quadratic selection functions are indicated by the thick faded blue (refugium) and green (native habitat) curves. The grey area called the “Death Plain” refers to a non-viable region (where a metapopulation with such a mean trait has negative growth rate at low density). In each subfigure, the mean trait at equilibria under changing environment at a given speed is indicated by a filled circle (resp. 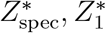 and 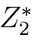). Notice how 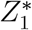 remains quite close to the initial state under stable environment 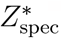 (Fig. 2b), while 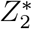 is far from it, closer to the refugium’s optimum than to the native habitat’s one (Section 2.3).

## 3 Results

Under a fixed environment (*c* = 0) and in the parameter range where a specialist species exists and is viable ([1 + 2*m <* 5*g*] ∧ [*g*(*m* − 1) *< m*^2^]), the analysis done in [Dekens 2022] shows that the selection gradient under stable environment ℱ^*c*=0^ cancels three times: in 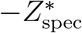, 0 and 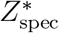, with respectively negative, positive and negative slopes (see Fig. 2a). This means that there exist three equilibria: two mirrored stable equilibria describing specialization to each habitat with respective mean traits 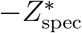 and 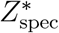, separated by an unstable equilibrium describing a generalist species (equally maladapted to both habitats). To understand the impact of a changing environment on the long-term adaptation of the focal species, I describe below the effect of increasing environmental speeds and increasing selection strengths on these equilibria. In my illustrations (Fig. 3, Fig. 4 and Fig. 5), I keep a constant intermediate migration rate for the sake of clarity (*m* = 0.5). I address sensitivity issues by displaying the analogous figures for a lower migration rate (*m* = 0.2) in Appendix G. Moreover, I test the results derived in this section with those from individual-based simulations in Appendix F (see in particular Fig. 6) to account for the influence of sampling effects and random demographic fluctuations .

**Figure 3.**
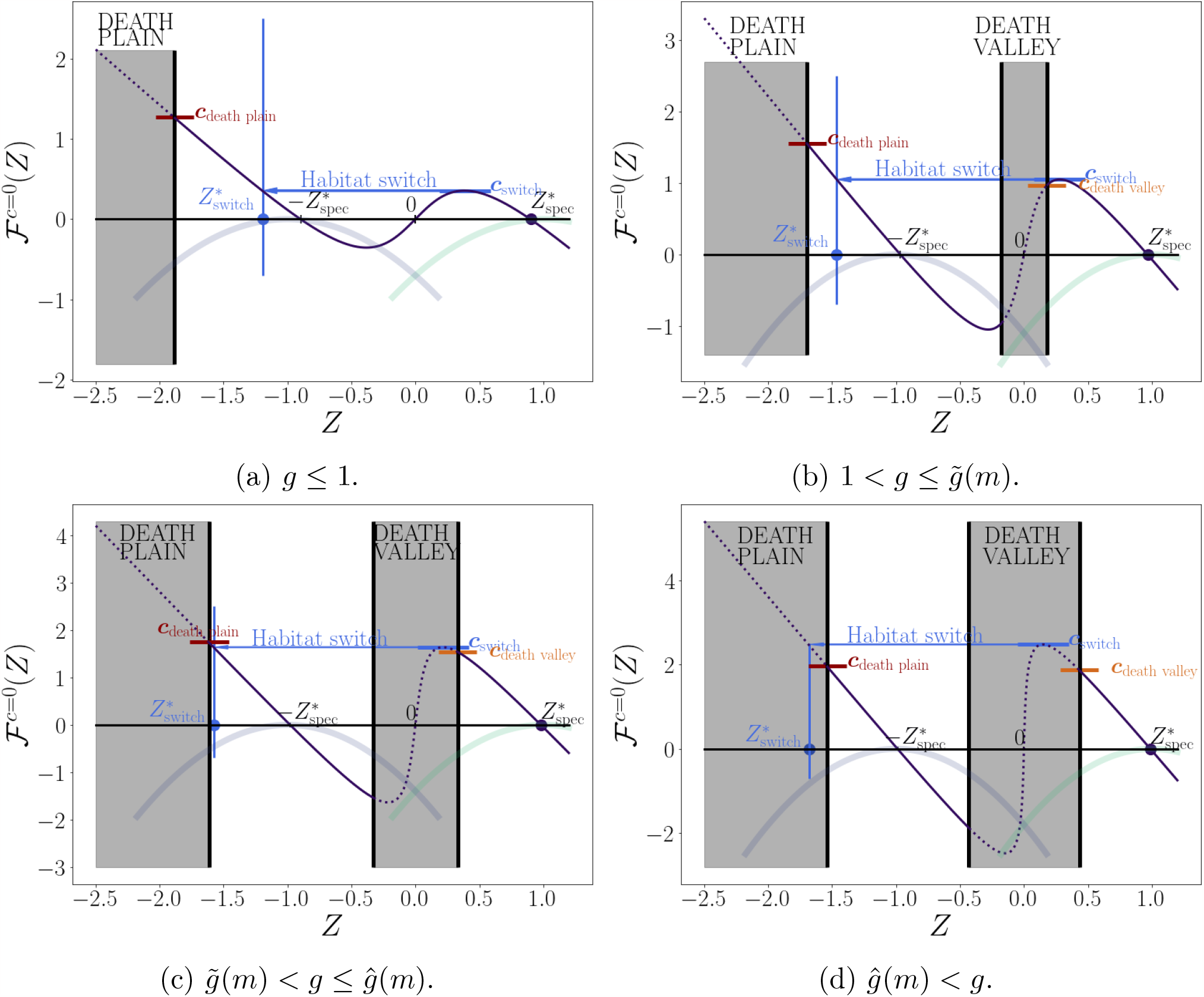
Critical transition speeds for increasing selection with *m* = 0.5. (from top to bottom, left to right, *g* = 0.7, 1.1, 1.4, 1.8). The tipping point provoking a habitat switch 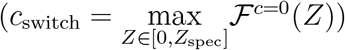 is indicated in blue, and the speeds corresponding to the non-viability area are indicated in orange (*c*_death valley_) and in crimson ((*c*_death plain_). Below *g* = 1, the death valley does not exist (Fig. 3a). Between 1 and 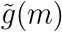, the switch occurs before entering the death valley (Fig. 3b). For 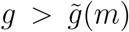, the switch occurs within the death valley. Below *ĝ*(*m*), the switch brings to a viable equilibrium (Fig. 3c) whereas it leads to extinction above (Fig. 3d).

**Figure 4.**
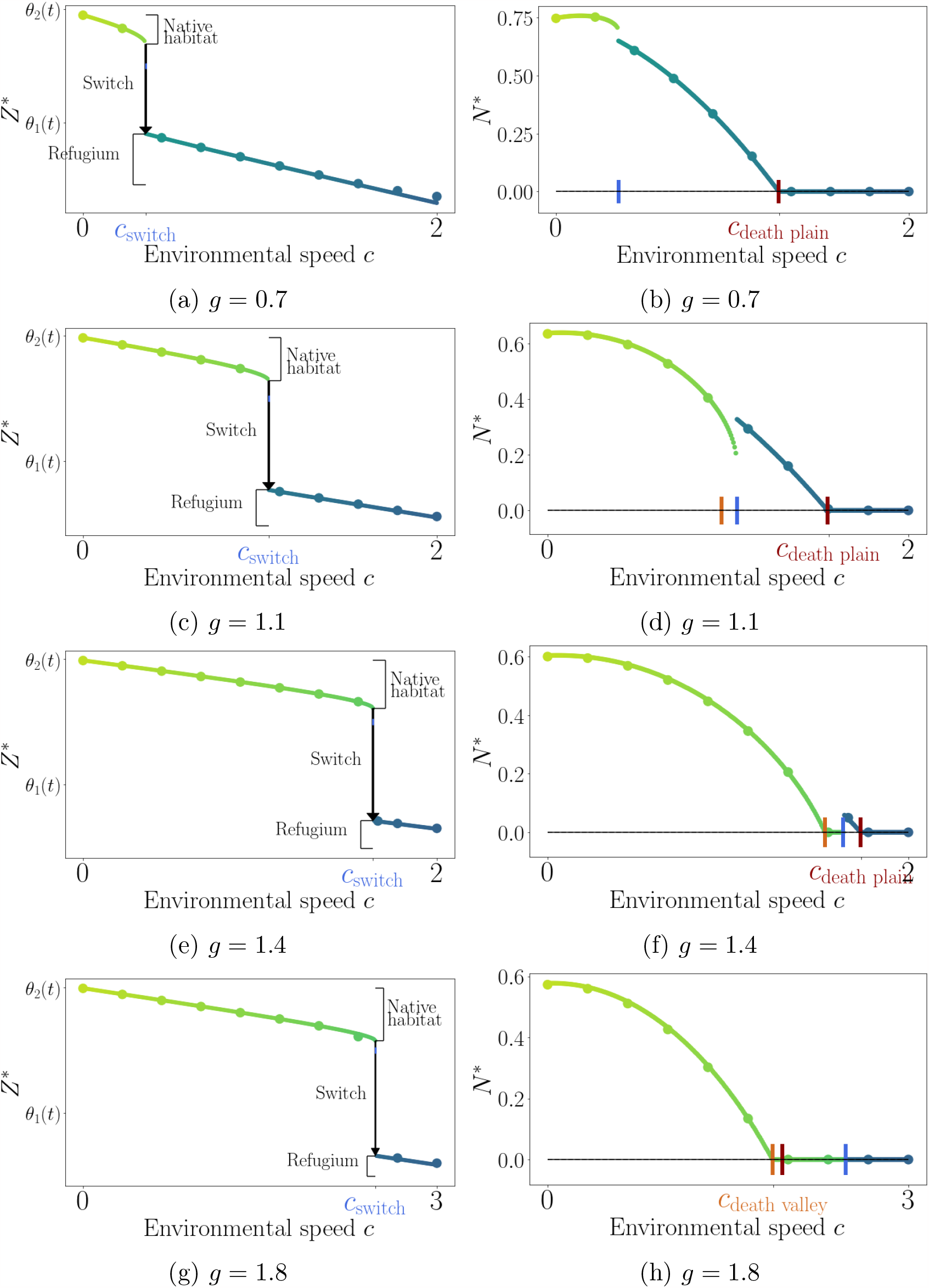
Equilibrium mean trait *Z*^*^ (left panel) and metapopulation size *N* ^*^ (right panel) as functions of the environmental speed *c* (*x*-axis), for increasing selection strengths corresponding to the ones used in Fig. 3 (top to bottom: *g* = 0.7, 1.1, 1.4, 1.8 ; with *m* = 0.5). Curves correspond to the analytical solution of Eq. (*S*) (green in the native habitat, blue in the refugium), and the dots are given by simulated numerical resolutions of Eq. (7). The vertical ticks in the left panel’s figures indicate *c*_switch_ (blue), *c*_death plain_ (crimson) and *c*_death valley_ (orange).

**Figure 5.**
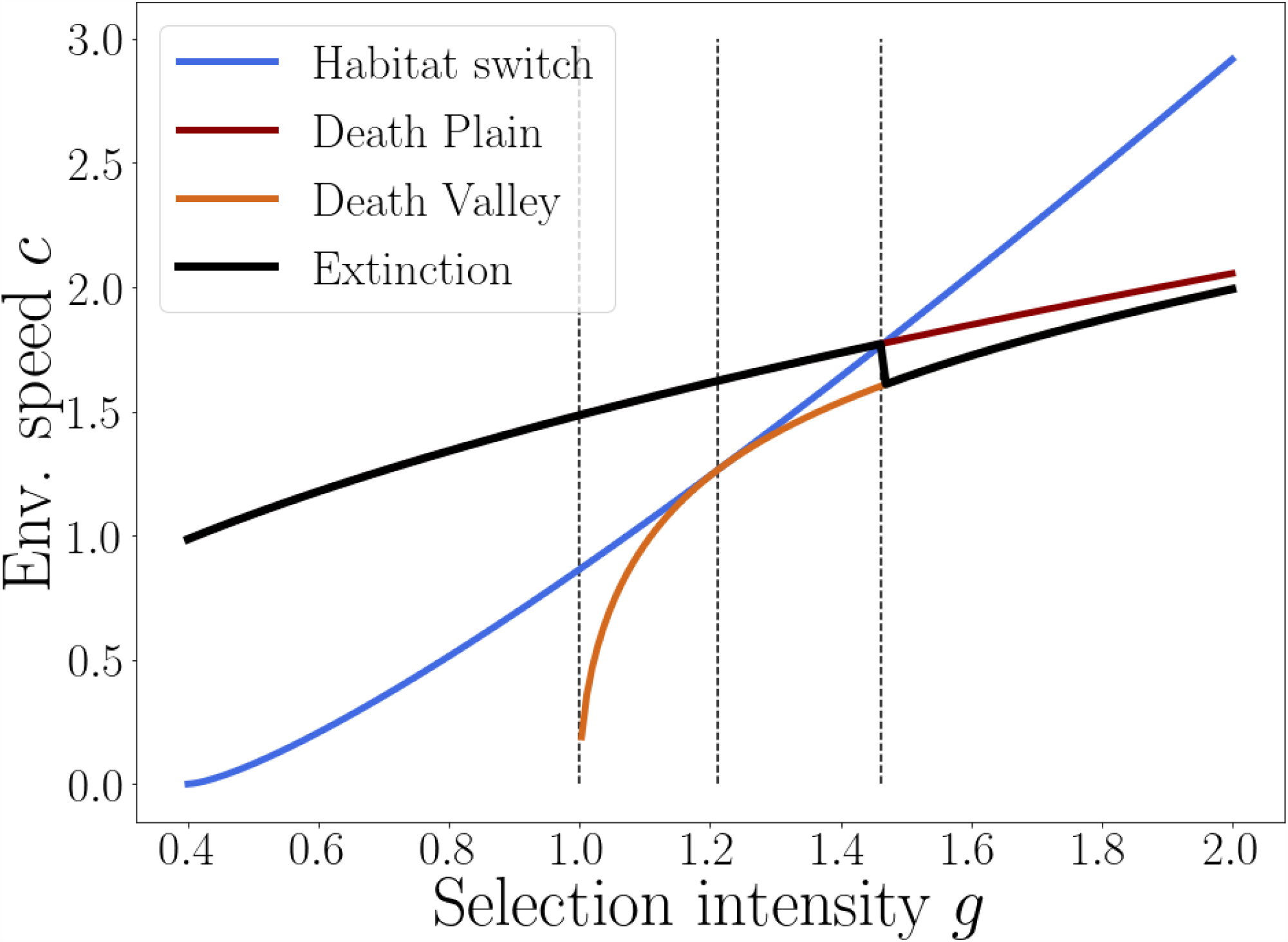
Analytical predictions of the critical speed of environmental change (black line) with increasing selection (*x*-axis), with *m* = 0.5. The coloured lines represent the different particular speeds that the analysis has identified and come from the analytical formula (3) for *c*_death valley_ and (5) for *c*_death plain_ and the identification of *c*_switch_ given in Section 3.2. The three vertical dotted lines delineate four selection regions corresponding to the ones illustrated in Fig. 3 and Fig. 4 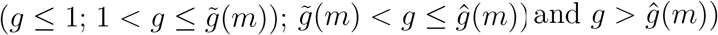.

### 3.1 Small environmental change: lagging behind the native habitat and beneficial loss of specialization?

For small speeds of environmental changes, the phase line indicates that the metapopulation’s mean trait at equilibrium *Z*^*^ lags behind the initial value 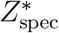 and thus increases its distance with the native habitat’s optimum (set at 1). This is reminiscent of the single-habitat models (see Kopp and Matuszewski 2014 for a review). However, on the contrary to these single-habitat models, where lagging is always deleterious for the population size, this can lead to an increase of the metapopulation size for small enough speeds of environmental change. In fact, I show in Appendix B that, regarding the transient dynamics at all environmental speeds, the metapopulation size actually increases immediately after the start of the environmental shift (at *t*≈0), as the mean trait *Z* starts lowering down from 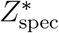. This initial increase in the metapopulation size occurs because, when the mean trait *Z*^*^ starts lagging behind its initial value, it actually becomes closer to the refugium’s optimum. Therefore, the immediate loss of population in the native habitat is overcompensated by the immediate gain of population in the refugium (even if it remains small), because the increase in adaptation for a trait far from an optimum is greater than the loss of it closer to an optimum when the selection functions are quadratic. This effect thus relies quite heavily on the precise shape of the selection functions, in particular their tails (for example, it does not occur when the selection functions are linear). This initial increase does not last, as, eventually, the lag with the native habitat increases and the increase of adaptation to the refugium is not enough to compensate the decrease of adaptation to the native habitat.

A related phenomenon is that, for very small speeds of environmental change, the metapopulation moves to an equilibrium 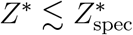 for which its size is actually greater that initially. The population as a whole is less specialized to the native habitat and benefits from its relative adaptation to the refugium.

### 3.2 Abrupt switch from the native habitat to the refugium: tipping point

In the last paragraph, we explained that, while for small environmental changes, the metapopulation can actually increase, it eventually starts dropping down as the lag increases. However, as opposed to the single-habitat model, the road to extinction here is not straight-forward, because of the existence of the refugium. I will first describe how this impacts the new equilibrium that is reached by the metapopulation, and next what can occur at the transient dynamics level on the path to this equilibrium.

#### New equilibrium

To determine the new equilibrium state reached by the population when the environment changes at speed *c*, one has to study the downward shifted phase line. As the phase line is smooth, it reaches a maximum value on [0, 1]. I call this maximum value 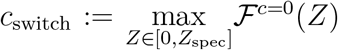, which is reached when *Z* = *Z*_switch_ *>* 0. This means that, for environmental change’s speeds that are lower *c*≤*c*_switch_, the metapopulation mean trait reaches an equilibrium value *Z*^*^ that is between *Z*_switch_ and 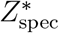 and thus closer from the native habitat’s optimum than the refugium’s one. Consequently, the metapopulation still remains in majority in the native habitat. However, for environmental change’s speeds that are even just greater than *c*_switch_, the metapopulation cannot reach an equilibrium with a positive mean trait *Z*^*^, as the phase line resulting from a downward translation does not cancel on [0, 1] anymore (see an example in Section 2.3). In our case, the only stable equilibrium that is left corresponds to a mean trait *Z*^*^ that is negative and even lower than 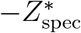, which is the mean trait of a metapopulation specialized to the refugium under a stable environment (*c* = 0). This means that *the metapopulation completely reverses its habitat preference* and now lags behind 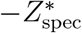, and becomes relatively better adapted to the refugium. *This shift from native habitat to refugium is abrupt*, because increasing the environmental speed even slightly above *c*_switch_ makes the possibility of remaining mainly in the native habitat completely disappear (see an illustration in Fig. 3a).

#### Tipping point

Moreover, once the metapopulation’s mean trait has dropped abruptly below 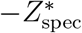, lowering the speed of environmental change back under *c*_switch_ (ie. translating the phase line slightly upward) does not result in the reversal of the habitat switch. Indeed, the metapopulation is now trapped in the basin of stability of specialization to the refugium. This indicates that the mean trait in the metapopulation has crossed a tipping point: for the metapopulation to become specialized in the native habitat again, the environment needs to actually change in the other direction, passing from a speed *c*_switch_ to a speed −*c*_switch_ (for symmetrical reasons).

#### Transient dynamics of the habitat switch

From the previous paragraphs, we know that the metapopulation will switch from mainly inhabiting the native habitat to the refugium when *c*≥*c*_switch_, because the new equilibrium is below 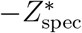. However, the latter does not describe how the switch occurs along the transient trajectory of the species. To do so, I will distinguish between two cases: intermediate selection 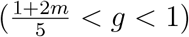 and strong selection (*g >* 1).

1. **Intermediate selection** 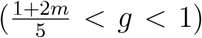: in this case, the analysis of viability done in Proposition 3.1 of [Dekens 2022] shows that the whole path that the mean trait *Z*(*t*) takes to go from lagging behind 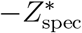 to lagging behind 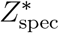 is viable. This means that for each 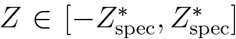, the metapopulation size *N* (*Z*) (defined through the ratio *ρ*(*Z*) satisfying the first line of *S*, see the details in B e.g.) is positive. This occurs because the selection is not strong enough for the local growth rates at low density to be both negative at any given mean trait of the metapopulation 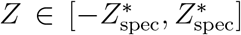. An example of this configuration is displayed in Fig. 3a.
2. **Strong selection** (*g >* 1): in this case, the analysis of viability done in Proposition 3.1 of [Dekens 2022] shows that the converse happens. The path from the native habitat’s optimum to the refugium crosses a non-viable stretch in the middle, that we call the *death valley*, which extends from −*Z*_death valley_ to *Z*_death valley_, where

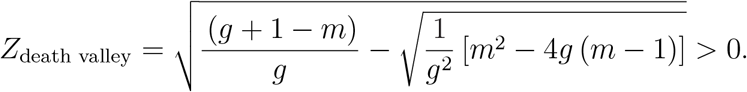

As the environmental change does not directly affect the subpopulation sizes, it also does not affect the boundaries of this death valley (see for example Fig. 3b). Concretely, when *Z*(*t*) enters the death valley, the metapopulation has a negative growth rate at low density and starts declining exponentially toward extinction. If the new equilibrium is in the death valley, the population goes extinct. However, if the mean trait *Z*(*t*) enters the death valley while going through a habitat switch/tipping point (for *c*≥*c*_switch_)), it means that, from the start of the transient trajectory, it is attracted by the new equilibrium *Z*^*^ beyond the death valley 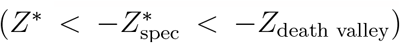. Thus, the switch occurs and forces the mean trait *Z*(*t*) to cross the death valley without stopping. In our deterministic model, this leads to dynamics of *evolutionary rescue* as the metapopulation manages to have its mean trait exiting the death valley before going extinct. The metapopulation then eventually bounces back primarily in the refugium. However, if selection is above a threshold depending on the migration rate denoted 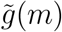, the possibility of switching between habitats occurs within the death valley (0 *< Z*_switch_ *< Z*_death valley_; see Fig. 3c). So, if the environmental speed *c* is such that *c*_death valley_ := ℱ^*c*=0^(*Z*_death valley_) ≤*c*≤*c*_switch_, the metapopulation is trapped at a stable equilibrium within the death valley without having the possibility to switch habitats, and goes extinct (see Fig. 4f for an illustration). The analytical expression for *c*_death valley_ is given by (see Appendix Appendix C for the derivation):

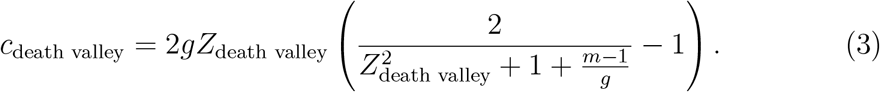

### 3.3 Lagging behind the refugium: connecting to the single-habitat models

From the previous section, if *c*≥*c*_*switch*_, the metapopulation switches from lagging behind the native habitat 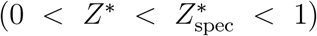 to lagging behind the refugium 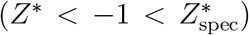. The two configurations are not symmetrical. In the former situation, the metapopulation’s mean trait is between the two local optima, so the situation is a compromise between adapting to the native habitat or the refugium. On the contrary, in the later situation, adaptation to the native habitat is really poor and the metapopulation relies entirely on the refugium. It is therefore reasonable to assume that the ratio between the subpopulation sizes reflect this: ie 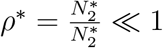 (this is hinted by the monotony of *Z*↦*ρ*(*Z*) with quadratic selection functions, see Appendix E). Using this in the right-hand side of the second line of *S* leads to an approximation of the lag of *Z*^*^ behind the refugium’s optimum (−1):

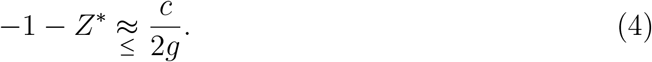

Simulations suggest that this approximation is quite accurate, as the native habitat’s influence on the metapopulation has almost completely faded, and the analysis now connects to the single-habitat framework, in which the lag is exactly (4) (note that in the timescale that is considered here, the population variance in trait is scaled to 1).

### 3.4 Critical speed for the metapopulation’s extinction

A crucial quantity in single-habitat models of adaptation to an environmental shift is the *critical speed for persistence*, below which the environmental changes slowly enough for the population to adapt and persist, and above which it changes too fast and the population goes extinct. In my two-habitats framework, persistence is not as clear-cut, as was pointed in the before-to-last section, where intermediate environmental speeds led to extinction. Instead, I use here *the critical speed for extinction of the population c*_extinct_ as the smallest speed such that ∀*c* ≥ *c*_extinct_, the population goes extinct at equilibrium. In this section, I explicit *c*_extinct_.

To do so, I rely once more on the analysis of viability under a stable environment in the Section 3.2 of [Dekens 2022]. It indicates that there exists a lower bound that I call *Z*_death plain_ below which the metapopulation’s mean trait *Z* always leads to negative growth rates at low density (leading to extinction), where:

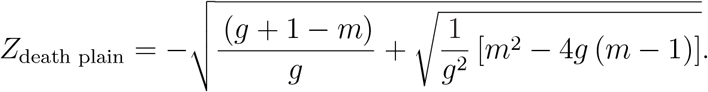

This lower bound *Z*_death plain_ *<* 0 defines a corresponding environmental speed *c*_death plain_ :=ℱ^*c*=0^(*Z*_death plain_) whose analytical expression reads (see a proof in C):

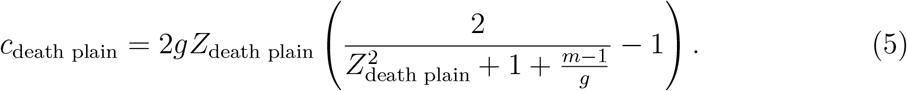

Moreover, there exists a selection threshold depending on the migration rate *ĝ*(*m*) *>* 1 such that *c*_switch_ ≤ *c*_death plain_ below and conversely above. Two cases can happen:

1. if *g* ≤ *ĝ*(*m*), for *c* ∈ [*c*_switch_, *c*_death plain_], the metapopulation switches to the refugium and lags behind the local optimum, but closely enough that it manages to persist (see Fig. 3a, Fig. 3b and Fig. 3c). If *c* ≥ *c*_death plain_, the lag in the refugium after the switch is too large and extinction occurs. <u>Therefore, in this case, the critical speed for extinction is *c*_death plain_</u> (see Fig. 4b, Fig. 4d and Fig. 4f). However, for some values of *g*, extinction can also occur for intermediate environmental speeds that are lower than the critical speed *c*_death plain_. Indeed, there exists a intermediate selection threshold 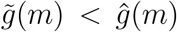 beyond which the switch occurs within the death valley (0 *< Z*_switch_ *< Z*_death valley_, see Fig. 3c). This means that for intermediate speed *c* ∈[*c*_death valley_, *c*_switch_], the metapopulation is trapped within the death valley without the possibility to switch, and goes extinct (see Fig. 4f).
2. if *g* ≥*ĝ*(*m*) *>* 1, we have *c*_death plain_ *< c*_switch_. Moreover, I show in Appendix D that, when *g* ≥1, the following inequality always holds: *c*_death valley_ *< c*_death plain_. Therefore, as <u>in this case the switch always occurs within the death valley (0 *< Z*_switch_ *< Z*_death valley_)</u>, for *c* ∈[*c*_death valley_, *c*_switch_], the metapopulation is stuck within the death valley without the opportunity to switch and goes extinct similarly as in the first case. However, in that case, when *c*≥*c*_switch_ and the switch toward the refugium does occur, it brings the population to a equilibrium where the lag with respect to the refugium’s optimum is too large and provokes extinction (see Fig. 3d). Therefore, in this case, the critical speed for extinction is *c*_death valley_ (see Fig. 4h).

From what came above, the critical speed of environmental change displays two key features in this two-patch framework . It is discontinuous at *g* = *ĝ*(*m*), because it passes from *c*_death plain_ (*g*≤*ĝ*(*m*)) to *c*_death valley_ (*g > ĝ*(*m*)), with *c*_death valley_ *< c*_death plain_. Consequently, it is also non-increasing with respect to increasing selection (see Fig. 5).

## 4 Discussion

### Summary

In this work, a two-patch quantitative genetic model with moving optima was used to analyse the eco-evolutionary dynamics of a sexually reproducing specialist species in a fragmented and changing environment comprising of its native habitat and a refugium. In a regime where the within-family variance is small, a separation of ecological and evolutionary time scales allows to reduce the complexity of the analysis to a phase-line study of the selection gradient. First, I show that small enough environmental speeds can actually be beneficial in terms of abundance thanks to a beneficial reduction of specialization. Moreover, for larger environmental speeds, there exists an evolutionary tipping point corresponding to a sharp habitat switch from the native habitat to the refugium. With strong selection, this shift brings the population to cross a death valley between the two habitats’ optima in the trait space, where the population size momentarily plummets and eventually rebounds when it reaches across, leading to evolutionary rescue. I finally compute the critical speed of environmental speed above which the population always goes extinct. This critical speed does not need to be increasing and can be discontinuous with respect to increasing selection strengths, especially with strong local selection.

### Critical speed of environmental change: fragmented spatial structure versus single-habitat

In Section 3.4, I derive the critical speed of environmental change above which the population goes extinct because it is too maladapted as a whole. Because of the fragmented nature of the two-patch environment available to the focal population and the existence of the refugium, this critical speed differs in a number of features as compared to the one classically derived in a single-habitat framework with quadratic selections (Lynch and Lande 1993; Bürger and Lynch 1995; Kopp and Matuszewski 2014).

First, here, the analysis shows that increased selection strength does not always equate to the population tracking the changing environment more efficiently, as it is the case in a single habitat with a quadratic selection function. More precisely, above a selection threshold, the critical speed of environmental change lowers down from *c*_death plain_ to *c*_death valley_ in a discontinuous fashion. This is due to the fact that with strong selection, the population can stay trapped within the rift between the habitats without the possibility to switch. A related result (even though it does not impact the critical speed of environmental speed) is that the population can go extinct for intermediate speeds of environmental change, which does not occur with a single-habitat framework.

Moreover, one could think that, because of the existence of the refugium, the population would be sheltered from extinction for larger environmental speeds than the critical speed derived in the analogous single-habitat models. Indeed, once the population has switched to lag behind the refugium, its adaptation is almost entirely determined by the dynamics within the refugium, as the migrants sent to the native habitat are too maladapted to make any significant contribution by gene flow. Therefore, the resulting dynamics are then very similar to the ones derived in the single-habitat framework, except for one major difference: the cost of dispersal. Due to the constant migration between habitats, there is a constant loss of population from the refugium that is not replenished. Thus, for a given lag in the refugium close to the critical one, the corresponding metapopulation size, approximated by the population size in the refugium, is actually smaller than without migration. Consequently, the critical environmental speed given by the present two-patch model is always lower than in single-habitat models.

### Fragmented spatial structure with moving optima versus to time-shifting environmental gradient in a continuous space

It would be tempting to use the present analysis as a literal stepping stone, by adding patches in a linear fashion to connect asymptotically with the continuous space models of adaptation to a time-shifting environmental gradient. However, I think that some cautions are needed when passing at the limit, because the space granularity is instrumental to obtain the qualitative results of this study, especially with strong selection. Indeed, with strong selection, a death valley of negative growth rate at low density appears between the two habitats, which is a key feature of fragmented environments whose translation in the continuous space limit is all but clear. Nevertheless, here, it underpins a significant part of the most original results, as the sharp dynamics of evolutionary rescue, but also the non-monotonicity of extinction (which can occur at lower environmental speeds than the critical one) and the discontinuous nature of the critical speed of environmental change with respect to increasing selection strength. The last two results differ from the results of quantitative genetics models considering an environmental gradient shifting at a constant speed in a continuous space setting (Pease, Lande, and Bull 1989; Polechová, Barton, and Marion 2009; Duputié et al. 2012; Aguilée et al. 2016; Alfaro, Berestycki, and Raoul 2017). Indeed, all of them conclude that there exists a critical speed under which the population always persist and beyond which it goes extinct. Note that there exists some variation due to the particular framework of each studies: for example, the mean trait speed can be lower than the environmental speed (which does not happen here) due to long range pollen dispersal (Aguilée et al. 2016) or in the case of a steep environmental gradient leading to a limited range equilibrium (Polechová, Barton, and Marion 2009). However, in both cases, it remains continuous with respect to increasing selection strength at the transition between qualitatively different equilibria.

That said, one can draw some qualitative parallels between this study and the framework of [Polechová, Barton, and Marion 2009]. Indeed, their authors show that increasing selection intensity (all else being held equal), therefore increasing the steepness of the environmental gradient, leads from a uniform range equilibrium (where the species invade the whole space and tracks the environmental gradient uniformly in space, with the same speed) to a limited range equilibrium (where the species only only significantly persists in a bounded spatial range and tracks the environmental gradient at a lower speed). This result is an extension of the one obtained in the study of [Kirkpatrick and Barton 1997] with stable environment (see also Mirrahimi and Raoul 2013; Raoul 2017). This echoes the passage from a generalist species equilibrium when selection is weak relative to migration (*g*≤1 + 2*m*) to a specialist species equilibrium when the converse holds (*g >* 1 + 2*m*) in the stable environment framework of [Dekens 2022] (note that, here, the study exclusively focuses on the parameter range *g >* 1 + 2*m* so on specialist species). However, as pointed out, the major difference is that the fragmented nature of the environment allows for sharp dynamics to occur that does not seem to exist in the continuous space setting. To reconcile the two frameworks will require additional technical work.

### Evolutionary tipping points in dynamics of structured populations

In Section 3.2, the analysis identifies tipping points in the dynamics of the population’s mean trait that lead to sharp habitat switches. When the environmental speed is below a threshold *c*_switch_, the population’s mean trait lags behind the native habitat’s optimum, but is closer from it than from the refugium’s optimum. Therefore, the bulk of the population is still located in the native habitat. (Just) Above the threshold, the population’s mean trait shifts abruptly to lag behind the refugium’s optimum, far from the native habitat’s one. Consequently, the population is suddenly relatively better adapted to the refugium. This sudden change of niche is difficult to reverse, because lowering down the environmental speed just below the threshold will not restore the initial configuration, as the population remains “trapped” in the refugium’s basin of stability. Restoring the population in its native habitat actually would require to completely reverse the environmental change, with the opposite speed (*c*_switch_→ −*c*_switch_.

Mathematically, one can visualize and predict such a tipping point thanks to the phase line study described in Section 3.2. A tipping point corresponds to a local maximum of the selection gradient (in our case the rightmost behind the initial mean trait). Assuming the selection gradient is smooth and non-constant, a sufficient analytical condition for the existence of a local maximum of the selection gradient is the co-existence of multiple equilibria (located where the selection gradient cancels). This is in agreement with what occurs in a single-habitat framework under changing environment with non-quadratic selection functions for which maladaptation stabilizes away from the optimum ([Osmond and Klausmeier 2017], [Garnier et al. 2022]). Indeed, in this case, the selection gradient under stable environment cancels in the optimum trait and converges to 0 in−∞. Heuristically, this means that the situation with an infinite lag is an asymptotic equilibrium. Therefore, between−∞and the optimal trait, the selection gradient reaches a local maximum, which corresponds to the evolutionary tipping point identified in the aforementioned studies. In these cases, past the tipping point, the lag grows indefinitely, so the population becomes extinct abruptly. In the present work, this does not occur, because the local selection functions are quadratic. Past the tipping point, the mean trait jumps on the stable branch of the selection gradient near the refugium’s optimum, so the population switches habitats abruptly and the lag stabilizes behind the refugium’s optimum. Moreover, here, either the jump brings the population to a viable state, or it was already extinct for environmental speeds just below the tipping point. So, in the present case, the evolutionary tipping point does not lead to an abrupt extinction, only to lagging behind the refugium’s optimum. However, this lack of extinction following a tipping point here might strongly be linked to the particular choice of quadratic selection functions.

To summarize, this work suggests that evolutionary tipping points can arise in stage-structured populations’ dynamics because of non-monotonic selection gradients, even with quadratic selection functions. This feature might be facilitated by naturally existing feedback loops between demography and evolution in these kind of models. Indeed, these lead to selection gradients towards integrative optimal traits that depend on the demographic state of the system, itself a function of the evolutionary state. For example, in the present work, this optimal trait is 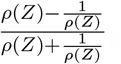, where 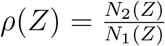 is the ratio between the two subpopulation sizes when the metapopulation mean trait is *Z*. A comparison with the analogous optimal trait obtained in the age-structured model [Cotto, Sandell, et al. 2019] suggests a correspondence between the elasticities quantifying the sensitivity of the population growth rate with respect to given transitions in their life cycle (introduced in Barfield, Holt, and Gomulkiewicz 2011) and the demographic terms*ρ*(*Z*) and 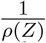 here. In our model, these quantities might thus be interpreted as being proportional to the elasticities linked to migration from one patch to the other. As such, they ponder the two local optima to build an integrative optimal trait for the global adaptation the metapopulation. Therefore, linking evolutionary tipping points and adaptation of stage-structure populations might come down to understanding the influence of such integrative optimal traits that account for different components of fitness on the co-existence of multiple equilibria in the system.

### Constrains in niche evolution and evolutionary rescue

The phenomenon of evolutionary rescue that is highlighted in Section 3.2 occurs because of the interplay between the sharp habitat switch and the existence of a death valley between the two local optima. This death valley is an area of the trait space (in our case, around *Z* = 0) where the growth rate at low density is negative and occurs when local stabilizing selection functions decline fast enough away from the optima. When the environmental speeds exceeds the threshold corresponding to the habitat switch, the still positive mean trait is attracted to an equilibrium with a negative mean trait. Therefore, during the habitat switch, it crosses the death valley. As soon as it enters it, the population size declines exponentially as the mean trait keeps lowering down to reach the lower bound of the death valley, where the growth rate at low density becomes positive again, which leads to a rebound of the population. This death valley connects with the concept of “fundamental niche limits” in moving optimum models highlighted in the review of [Klausmeier et al. 2020]. In a changing environment, it might happen that the species adapt not only by shifting its spatial distribution, but also by shifting its niche, like in [Aguilée et al. 2016]. However, it might happen that the niche is also biologically constrained, defining regions of positive and negative growth rate at low density (the latter in our case comprising the death valley). If the adaptive trajectory crosses such a region, the population size plummets, but can be saved if if comes back into the fundamental niche (see Fig. 6 of Klausmeier et al. 2020 for an illustration). It is noteworthy to point out that this phenomenon of evolutionary rescue, as in our case, does not rely on the advent of beneficial de novo mutations, rather on standing genetic variation within the population prior to entering the death valley. This standing genetic variation is due to the redundancy of the highly polygenic genetic architecture with numerous small effects that shift in frequency along the adaptive trajectory and segregates due to sexual reproduction. Moreover, the model I use here does not allow to directly study the trajectory of the population within the death valley. In particular, the analysis does not allow to quantify the speed with which the mean trait crosses the death valley versus the decline of the population, which would inform on the probability of rescue. In fact, since the framework used here is deterministic, rescue always occurs and the metapopulation carries on, sometimes reaching a non-extinct equilibrium.

### Limits of the model

The last paragraph introduces some of the limits of the model presented in this paper. Although its deterministic features allow both to work with clear assumptions in a well-defined framework and to derive clear-cut results, they are not equipped to capture finer phenomena, in particular the ones where stochasticity is central. To account for the influence of sampling effects and random demographic fluctuations, I performed supplementary individual-based simulations (IBS) described in Appendix F and compare their results with the analytical predictions of my model (see Fig. 6). The first conclusion is that the results of the IBS are in excellent agreement with the analytical predictions of the present model whenever the equilibrium state of the population is either far from extinction, or definitely extinct. However, as the IBS are subject to random demographic fluctuations, they give a more nuanced conclusion when my model predicts that the population gets rescued to get through the death valley. In fact, they show that the metapopulation does not get rescued in a significant part of replicate simulations. However, in some replicate simulations, it does, and then the IBS result in an equilibrium that is very close to the one my analysis predicted (see Fig. 6d). The IBS give an estimation of the probability of rescue (which cannot be quantified by my approach) and put in perspective the analytical predictions derived here: the latter describe the equilibrium well, conditional on the metapopulation getting rescued.

To take matters even further than accounting for the influence of demographic stochasticity and sampling effects, one might focus on the central hypothesis of this model, the one about the within-family segregational variance. A key feature that makes the model analytically tractable is that it is fixed, therefore summarizing in one parameter all the details of the underlying genetic architecture (the IBS I performed make the same assumption in order to isolate the influence of population dynamics’ stochasticity). A first direct consequence is that this framework does not account for the biological constraints that a finite genetic architecture imposes on the possible phenotypic range. The latter would be necessarily bounded and will be exceeded in the long run by a never-ending shifting optimum, making it impossible for the population to adapt without de novo mutations (connecting once again with the fundamental niche limits of Klausmeier et al. 2020). Furthermore, it is known that in a finite population in a stable environment, this segregational variance becomes eroded by inbreeding through time (see Barton, Etheridge, and Véber 2017 for a quantification of the error that builds up). This is presumably enhanced during rescue events, where the population goes through a significant bottleneck, which is known to reduce the within-family genetic variance severely (or conversely, increase the probability of identity). Whereas in our model, the population picks up after an episode of rescue with the same segregational variance, allowing it to persist in the long run in some cases by stabilizing its lag with the refugium’s optimum, I presume that stochastic individual-based simulations with explicit genetic architecture would show that an *extinction debt* can accumulate as a result of maladaptation and the loss of genetic variance, dooming the population quite soon after the rebound (relating to the concept introduced in Tilman et al. 1994). To quantify both the probability of evolutionary rescue and the long-term consequences of the loss of genetic variance would require to adopt a stochastic modellign approach with an explicit genetic architecture.

### Perspectives

As detailed above, a modelling choice that harbours the potential to significantly change the qualitative results of moving optimum quantitative genetic models is the one about the (local) selection function. Indeed, for example, in the single-habitat framework, [Osmond and Klausmeier 2017], Klausmeier et al. 2020 and Garnier et al. 2022 highlight how choosing a selection function where maladaptation stabilizes away from the optimum instead of quadratic selection functions where maladaptation increases faster and faster away from the optimum leads to the occurrence of evolutionary tipping points. One can thus wonder how making a similar choices would alter the results here and how to analyze a model with different selection functions. As a matter of fact, the analytical steps used in this study (justification of the Gaussian approximation of local trait distributions under small within-family variance and separation of ecological and evolutionary time scales leading to the phase line study) are robust to using other selection functions: I focused here on the quadratic case here because it allows to derive explicit analytical results. With another choice of selection function, the precise expression and properties of the phase line linked to the selection gradient will depend on the particular choice and potentially lead to qualitatively different results. However, the general method used here for analyzing the resulting phase line is also transferable to another one arising from a different selection gradient.

Besides, the major hypothesis underpinning the analysis and therefore the results is that the within-family variance due to the segregation of the many small-effect loci underlying the adaptive quantitative trait is small compared to the distance between the local optima (but also more generally all the other parameters). This regime has been the analytical frame of several studies modelling the trait inheritance with the infinitesimal model (Calvez, Garnier, and Patout 2019; Patout 2020; Garnier et al. 2022; Dekens 2022). It provides sufficient standing genetic variance for selection to act upon to shift the population’s mean trait, albeit on a slower time scale than the ecological dynamics governed by birth, death and migration processes, which allows one to justify the Gaussian approximation of local trait distribution that is classical for quantitative genetic models. As this hypothesis leads to the sharp dynamics presented in Section 3, one could ask what would happen if one relaxes this assumption. Beyond a purely theoretical question, this could have concrete implications for conservation purposes: in the case of niche specialist species facing climate change, would it beneficial to try and increase the standing genetic variance to promote local adaptation and would it make it more resilient to greater environmental speeds? A further study is required to address this issue.

## Data availability

The codes to reproduce the figures of this artile are available at https://github.com/ldekens/two-patch-model-changing-environment.

## Acknowledgements

I first thank Vincent Calvez for reviewing this manuscript and offering great suggestions to improve it, and Ophélie Ronce for initially introducing me to the biological question question underlying this work and for subsequent insightful discussions. I also thank Sepideh Mirrahimi, Amandine Véber and Sally Otto for valuable feedback. I thank the Foundation for Mathematical Sciences in Paris (FSMP) for funding my work and acknowledge having received partial funding from the ANR project DEEV ANR-20-CE40-0011-01 and from the European Research Council (ERC) under the European Union’s Horizon 2020 research and innovation program (grant agreement No 865711).

## A Detailed derivation of the limit system *S*

**Dimensionless system** In the regime where the within-family variance ***σ***^2^ is small compared to the difference between the local optima (***θ***_**2**_−***θ***_**1**_ = 2***θ***), it is convenient to define the following rescaled variables and parameters to get a dimensionless system from (2):

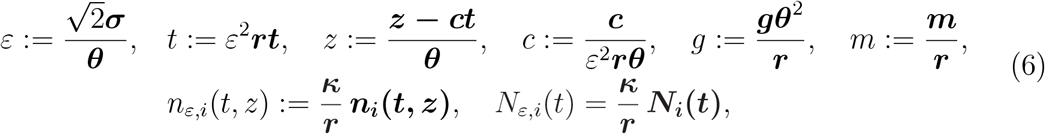

and the infinitesimal model reproduction operator ℬ_*ε*_[*n*_*ε,i*_](*t, z*) =ℬ_***σ***_**[*n***_***i***_**](*t, z*)**. Rescaling (2) according to (6) yields the following system:

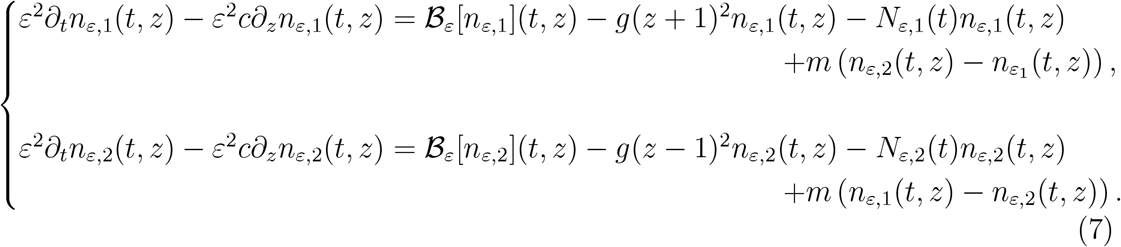

The change in the trait variable 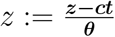 means that we place ourselves in the moving-window frame which moves at the same speed as the environment. In this referential, the local optima are fixed, but the environmental shift’s action appears in an additional advection term (where the factor *ε*^2^ comes from the change in the time variable):

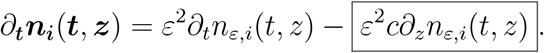

### Moment-based system in the regime of small within-family variance

In the regime of small within-family variance, [Dekens 2022] showed that the moment-based system obtained from integrating (7) is closed, as the trait distributions *n*_*ε*,1_ and *n*_*ε*,2_ are approximately Gaussian, of small variance 2*ε*^2^. Its derivation is similar as in [Dekens 2022], except for the advection term −*c∂*_*z*_*n*_*ε,i*_ due to the changing environment, whose integration I detail below.

As the local distributions *n*_*ε,i*_ are expected to stay concentrated, the speed of the environmental change does not impact directly the dynamics of the subpopulations sizes at the main order, as ∫_ℝ_ *∂*_*z*_*n*_*ε,i*_(*t, z*)*dz* ≈ 0. However, it does impact directly the dynamics of the local mean traits, since one can integrate by parts to obtain

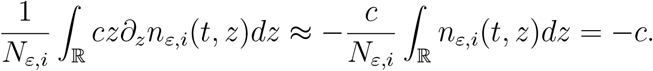

Due to the last computation, the moment-based system obtained from integrating (7) here yields

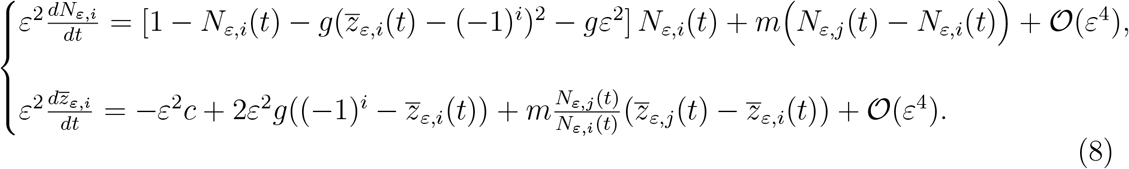

### Separation of time scales

Introducing the same slow-fast variables 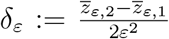 (trait discrepancy between habitats) and 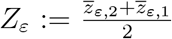 (average trait in the metapopulation) and denoting by *ρ*_*ε*_ *>* 0 the ratio between subpopulation sizes: 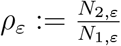 as in [Dekens 2022], one can obtain the following slow-fast system from (8):

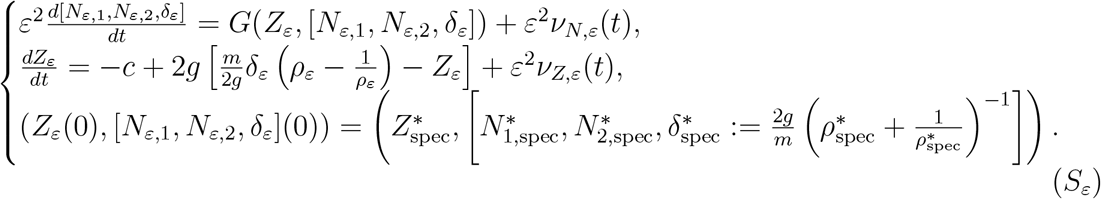

The function *G* and the residues *v*_*N,ε*_ and *v*_*Z,ε*_ are defined identically as in [Dekens 2022] (Eq. 18).

#### Remark A.1

(Direct influence of the environmental change in *S*_*ε*_). *The fast dynamics of the subpopulation sizes encoded by G (first line of* (*S*_*ε*_)*) are independent from the environmental shift, because the latter only directly impacts the dynamics on the local mean traits in* (8) *(second line). This results in the term* −*c in the slow dynamics on the average trait Z*_*ε*_ *(second line of* (*S*_*ε*_)).

#### Remark A.2

(Initial conditions: asymmetrical equilibrium for specialists of habitat 2.). *The initial state of the system* 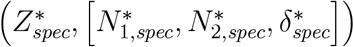 *has two particularities that follow its definition in Proposition 4*.*2 in [Dekens 2022]. First, it describes a specialist species that is mainly adapted to the native habitat (*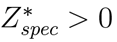 *is close to its local optimum) and mainly inhabits this habitat (*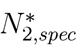 *is much larger than* 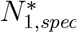, *which means that* 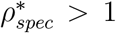*). Second, the initial specialist population hereby described is at equilibrium under a stable environment* (*c* = 0). *This means that it both cancels the fast dynamics of the subpopulation sizes represented by the function G in S*_*ε*_:

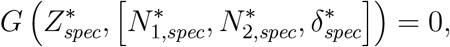

*and the slow dynamics of the average trait Z*_*ε*_ *(right-hand side of the second equation of S*_*ε*_ *at the main order when c* = 0):

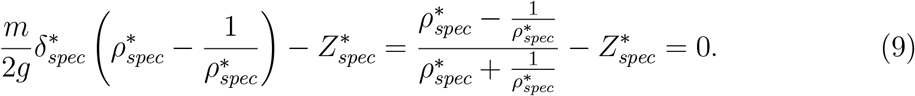

The two previous remarks justifies the separation of time scales, as it relies on the local stability of the fast dynamics, which is the same as in [Dekens 2022]. Therefore, the time scales between demographic dynamics and trait dynamics can be separated when *ε*→0, as stated by Theorem 3.1 of [Dekens 2022], whereby the solutions of (*S*_*ε*_) converge to the solutions (*Z*(*t*), [*N*_1_(*t*), *N*_2_(*t*), *δ*(*t*)]) of the following system:

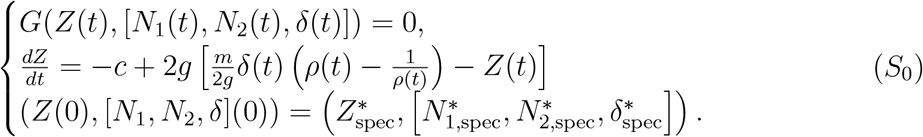

There are two advantages to *S*_0_. First, due to the separation of time scales, the de-mographical dynamics are instantly resolved for any current value of the average trait in the metapopulation *Z*(*t*) (as the first line of *S*_0_ is an algebraic equation, not a differential one). Second, the metapopulation is monomorphic, which is revealed by the fact that the dynamical variable *Z*(*t*) is the average trait in the metapopulation. This is because the gene flow by migration occurs at the fast time scale, as opposed to the shift of local mean traits due to selection. As a result, the two local mean traits merge on the fast time scale into *Z*(*t*), which then moves slowly according to the gradient of selection represented by the right-hand side of the second line of . More precisely, the latter pushes toward an integrative optimum 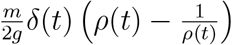 resulting from the demographical balance of the system.

Moreover, the analysis done in [Dekens 2022] allows to further simplify the system *S*_0_. It shows that the population can actually be fully described by its average mean trait *Z* and the ratio between subpopulations sizes 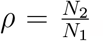 . Their dynamics are given by *S*.

## B Metapopulation size’s initial increase

In this appendix, I show that 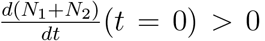, which implies that, when the environment starts changing, the metapopulation size increases. The intuition is that at the start of environmental change, the initially deserted refugium gets more suited to support the species as the native habitat becomes less suited. However, because of the quadratic selections functions, the gain of quality in the refugium is greater (initial selection gradient of 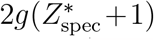 that the loss of quality in the native one (initial selection gradient of 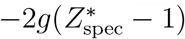.

*Proof*. Following the notations of Lemma 1 in Dekens 2022, I define 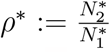 the ratio of subpopulation sizes and the following polynomial form: *P*_*z*_(*X*) :=*X*^3^ − *f*_1_(*z*)*X*^2^ +*f*_2_(*z*)*X* − 1, with 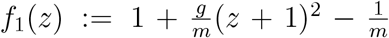 and 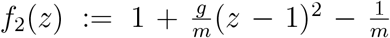 .

From Lemma 1 in [Dekens 2022], for *Z*^*^ℝ, 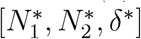 is a fast equilibrium at the level 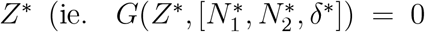 with 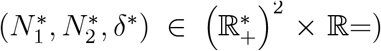 if and only if 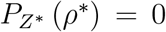and *ρ*^*^ *>* max(0, *f*_1_(*Z*^*^)). In this case, (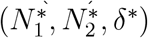 is given by 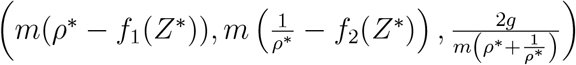.

The system (*S*_0_) is therefore equivalent to

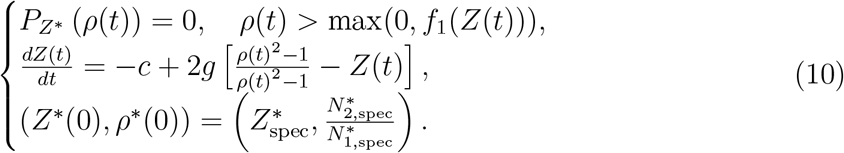

Hence, the derivative of the metapopulation size à time *t* = 0 is given by

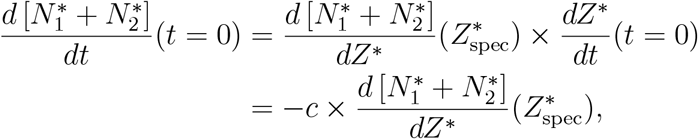

thanks to (9).

It is therefore sufficient to show that 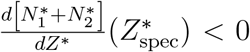. Recalling that 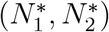 is given by *m*(*ρ*^*^ − *f*_1_(*Z*^*^)), 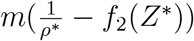) where 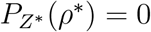, we deduce that:

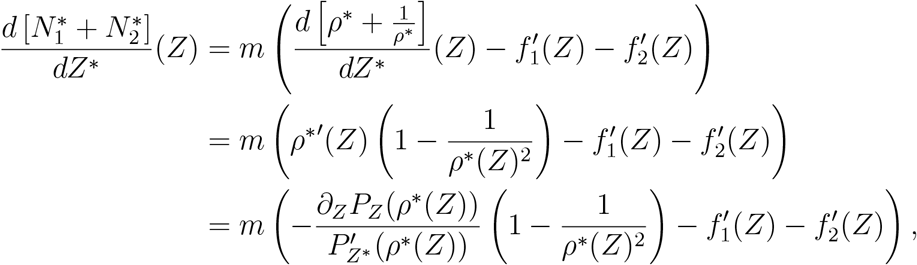

where I used that 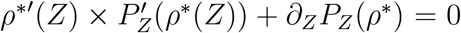 (since *P*_*Z*_(*ρ*^*^(*Z*)) = 0) and that 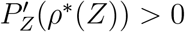, since *ρ*^*^(*Z*) is the largest root of *P*_*Z*_ (and has multiplicity one).

Moreover, because *P*_*Z*_(*X*) = *X*^3^ − *f*_1_(*Z*)*X*^2^ + *f*_2_(*Z*)*X* − 1 with and 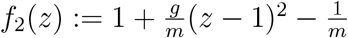, we obtain that

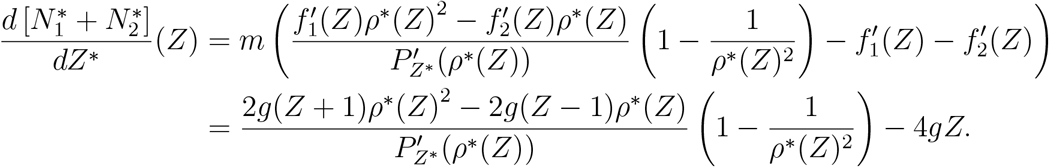

Since the computation is perfomed at *Z* = *Z*_spec_ that satisfies 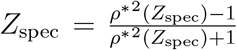,we obtain that

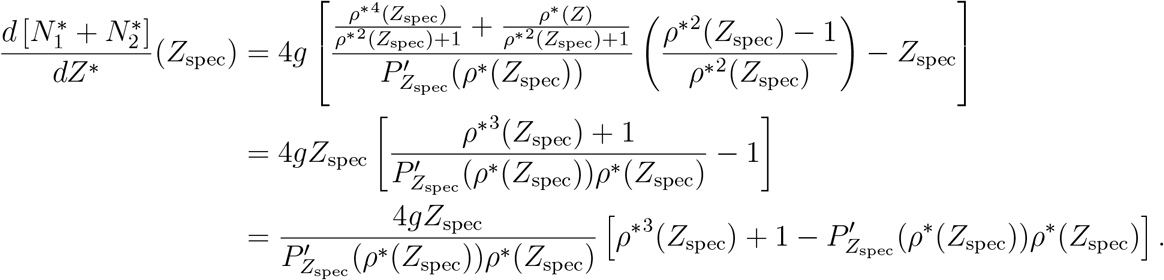

Since *Z*_spec_*>*0 by hypothesis, that 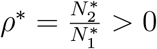 is the largest root of *P*_*Z*_ with multiplicity one (see lemma 2 of Dekens 2022) and that therefore 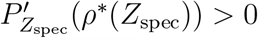, it is sufficient to determine the sign of the term within brackets to conclude:

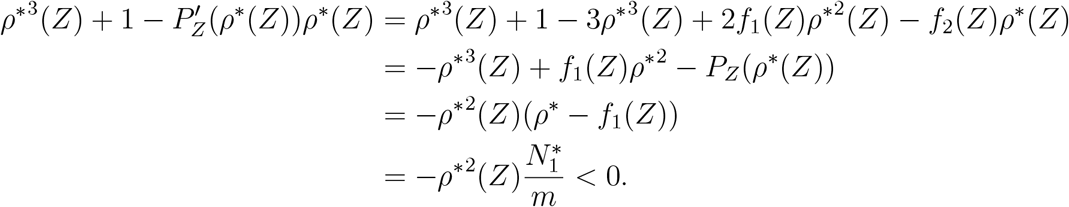

Hence the result.

## C Derivation of *c*death valley and *c*death plain

Explicit analytical expressions for *c*_death valley_ and *c*_death plain_ are available in the parameter range of interest ([1 + 2*m <* 5*g*] ∩ [*m*^2^ *>* 4*g*(*m* − 1)]). One can compute that:

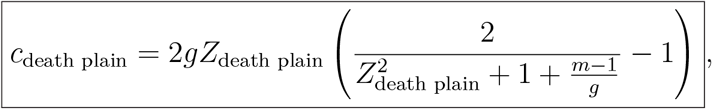

With

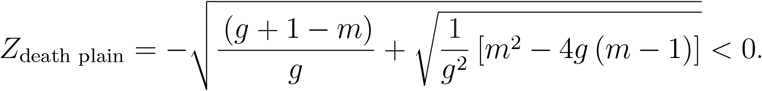

Similarly, when *g* > 1:

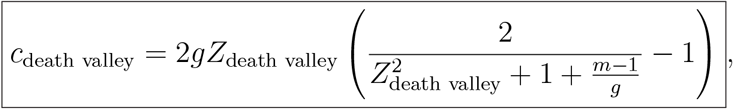

With

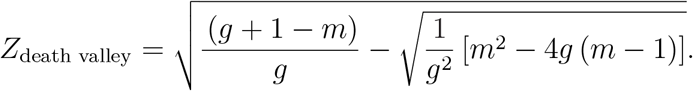

*Proof*. At a dominant trait *Z*^*^, the fast equilibria are defined by 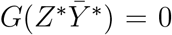, which is solved by (Lemma 1, Section 3.2 of [Dekens 2022])

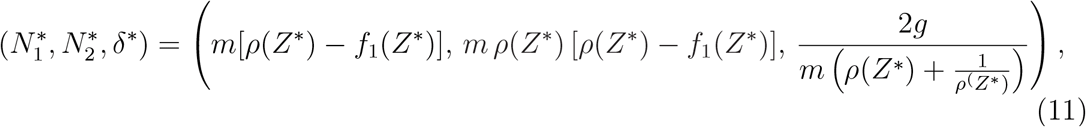

where *ρ*(*Z*^*^) is the largest (positive root) of the cubic polynomial

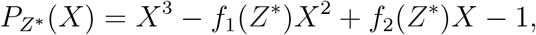

and

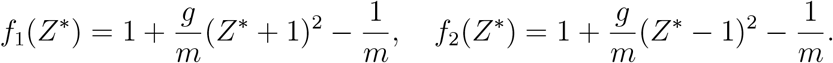

(11) implies that the subpopulations sizes 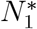 and 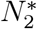 are positive if and only if *f*_1_(*Z*^*^) *< ρ*(*Z*^*^). Therefore, the limit of viability occurs for *Z*^*^ such that *f*_1_(*Z*^*^) = *ρ*(*Z*^*^). As *ρ*(*Z*^*^) is the largest root of 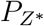, the latter leads to 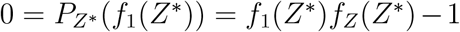, whose roots are ±*Z*_death plain_ and ±*Z*_death valley_. Consequently, the lowest limit of viability occurs at *Z*^*^ = *Z*_death plain_ *<* 0. The corresponding critical speed over which the population goes extinct reads (renaming *Z*_death plain_ as *Z*_*DP*_ for the computation)

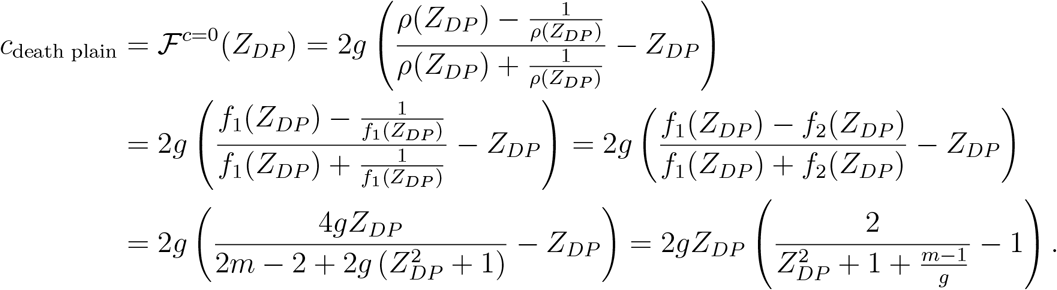

The same applies to *c*_death valley_ and *Z*_death valley_.

## D Proof that *c*_death valley_ *< c*_death plain_ for all *g >* 1

Suppose that *g >* 1 in addition to the parameter range of interest ([1 +2*m <* 5*g*] ∩ [*m*^2^ *>* 4*g*(*m*−1)]). I rename here *Z*_death plain_ and *Z*_death valley_ as *Z*_*DP*_ and *Z*_*DV*_ respectively, for the length of this proof. I recall that the limit traits for the viability of the population *Z*_*DV*_ *>* 0 *> Z*_*DP*_ are defined by their squares being the two real roots of the polynomial *X*^2^ −*SX* +*P*, with 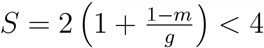 and 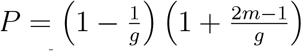 (in the considered parameter range, Proposition 3.2 in [Dekens 2022] ensures that the discriminant *S*^2^ −4*P* is positive). Moreover, standard algebra implies that 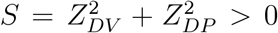 and 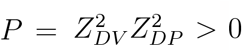. For the sake of clarity, I gather all the inequalities that will be used in the rest of the proof.

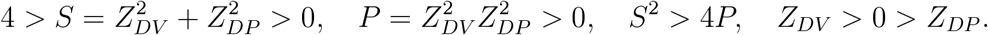

Using the expression of *c*_death plain_ and *c*_death valley_ derived in Appendix C, one can compute (defining 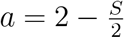):

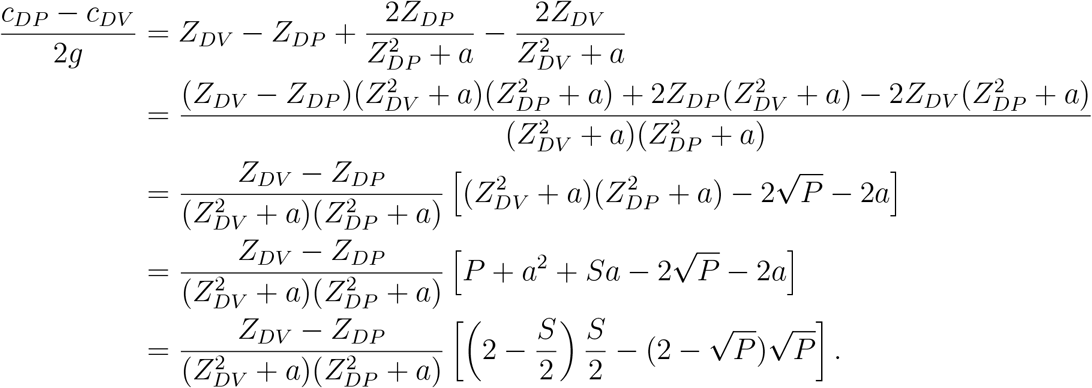

Since *Z*_*DV*_ *>* 0 *> Z*_*DP*_, *c*_*DP*_ − *c*_*DV*_ has the sign of 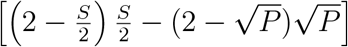 . Defining the function *f* : *x* ↦ *x*(2 − *x*), it remains to show that 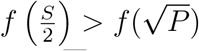.

However, on the one hand, from the inequalities above,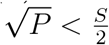. On the other hand, from the expression of *P* and *S*, one can compute

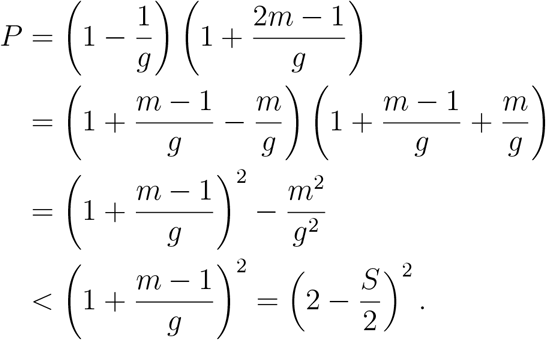

As 0 *< S <* 4, we obtain that 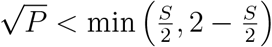 . Hence, since *f* is strictly increasing on [0, 1], that either 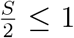 or 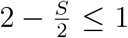, and that 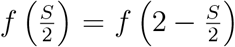, we deduce that 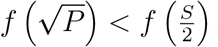, which concludes the proof.

## E Monotony of *ρ*(*z*)

Let *z* be such that *ρ*(*z*) is the only positive root of *P*_*z*_(*X*) = *X*^3^−*f*_1_(*z*)*X*^2^ + *f*_2_(*z*)*X*−1 greater than *f*_1_(*z*), so that viability is ensured (see Dekens 2022 for details). From *P*_*z*_(*ρ*(*z*)) = 0, we get

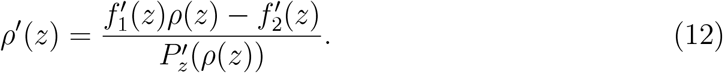

As *ρ*(*z*) is the largest root of *P*_*z*_ (see Dekens 2022), 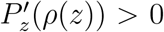. Therefore, *ρ*^′^(*z*) has the sign of 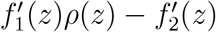.

Example of monotony: in the case where *f*_1_ is increasing on [−1, +∞[ and *f*_2_ is decreasing on ] − ∞, 1] (like in the quadratic case), then *ρ* is an increasing function of *z* ∈ [−1, 1].

## F Supplementary figures from stochastic IBS’s results

In this appendix, I compare the analytical predictions regarding the equilibrium values of the metapopulation size and mean trait derived in Section 3 with the end results of stochastic individual-based simulations (IBS) with discrete and overlapping generations. The migration and selection parameters used in the following display are those used in Fig. 4, which are *m* = 0.5, *g* = 0.7, 1.1, 1.4, 1.8.

### IBS’s design

The IBS are designed as follows. For a given selection parameter *g, N*_replicates_ replicate simulations are run for a fixed number of generations *N*_gen_ that is sufficient to reach the equilibrium, after 100 generations of burn-in. Moreover, each generation spans a small time-step *dt* = 10^−2^ (so that only a few events occur at each generation). For *g* = 0.7, 1.1, 1.4, 1.8, *N*_replicates_ = 250, 1000, 1000, 1000 and *N*_gen_ =3.5×10^6^, 2×10^6^, 10^6^, 5×10^5^. The large number of replicates meant to ensure that (rare) evolutionary rescue events would be captured in most cases. Notice that the number of replicates for *g* = 0.7 is relatively low (250, which is still quite large), as the analysis predicts that no evolutionary rescue is expected in this case and that cutting the number of replicate simulations speeds up the computational time. Each replicate simulation is run according to the following:

⋄ Each habitat has the same carrying capacity *K* = 10^4^ individuals. Initially, there are 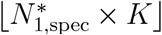 individuals in the refugium and 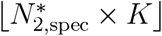 individuals in the native habitat, where 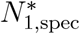 and 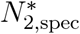 are the rescaled equilibrium subpopulations sizes under stable environment indicated in Proposition 4.2 of [Dekens 2022]. Their traits are randomly drawn from a Gaussian distribution of mean 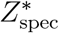 and with a small variance *ε*^2^, with *ε*^2^ = 5 × 10^−3^.
⋄ First, the environment is stable during 100 generations. Next, the environment changes with a speed *ε*^*2*^*c*, meaning that in habitat *i* and at generation *t* after the burn-in, the local optimal trait is given by *θ*_*i*_(*t*) = *θ*_*i*_(0) + *ε*^2^*ct*, with 1 ≤ *t* ≤ *N*_gen_.
⋄ At each generation *t*, the following life cycle happens:
  1. Reproduction event: in each subpopulation *i*, a random number of individuals are uniformly sampled across the subpopulation (*N*_*i*_(*t*) ×*dt* individuals chosen on average). For each of these individuals, a mate is uniformly chosen at random within the same population. Their mating produces a child added to the subpopulation and whose trait is drawn randomly from a Gaussian distribution centered on the mean parental trait and with variance 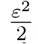. At the end of the reproduction event, the subpopulation size is 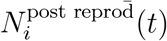.
  2. Selection-competition event: in each subpopulation living in habitat *i* with optimal trait *θ*_*i*_(*t*), each individual faces a selection-competition trial, according to its trait *z* and the current subpopulation size 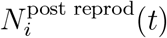. Precisely, individuals survive according to independent Bernoulli random variables with parameters given by:

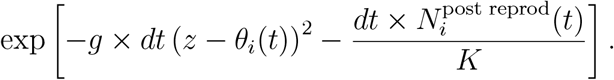

Any individual who fails the trial is removed and the subpopulation size at the end of this phase is 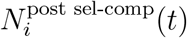.
  3. Migration event: in each subpopulation *i* independently, a random number of migrants is drawn according to a Poisson distribution of parameter 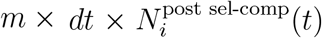. These migrants are then removed from their current subpopulation and added to the other one.

### IBS’s results

The comparison between the analytical predictions of the equilibrium variables (*Z*^*^ the metapopulation mean trait and *N* ^*^ the metapopulation size) and the analogous final quantities of the IBS is displyed in Fig. 6. The overall conclusion is that the IBS are in excellent agreement with the predictions, especially for the metapopulation mean trait *Z*^*^ conditional on persistence (left column). The same can be said for the metapopulation size, albeit with more variance near extinction (vertical black lines, representing 98% of the trajectories) as expected by random stochastic fluctuations. These have a striking impact when the metapopulation relies on evolutionary rescue, which is predicted to occur when *g >* 1 (bottom three lines), and is highlighted particularly in Fig. 6d. Notice that for the 6th, 7th and 8th environmental speeds, the median metapopulation size (black squares) is 0, indicating extinction. However, the variance between replicates can be high and the median metapopulation size of non-extinct populations (black circles) is very close to the analytically predicted metapopulation size. The IBS give some quantitative indications that the probability of rescue is probably less than half and more than 1*/N*_replicates_ = 10^−3^ (whereas this probability of rescue cannot be quantified from my analysis). Moreover, it shows that when the population does get rescued, its final state is accurately predicted by my analysis.

## G Supplementary figures with low migration rate *m* = 0.2

**Figure 6.**
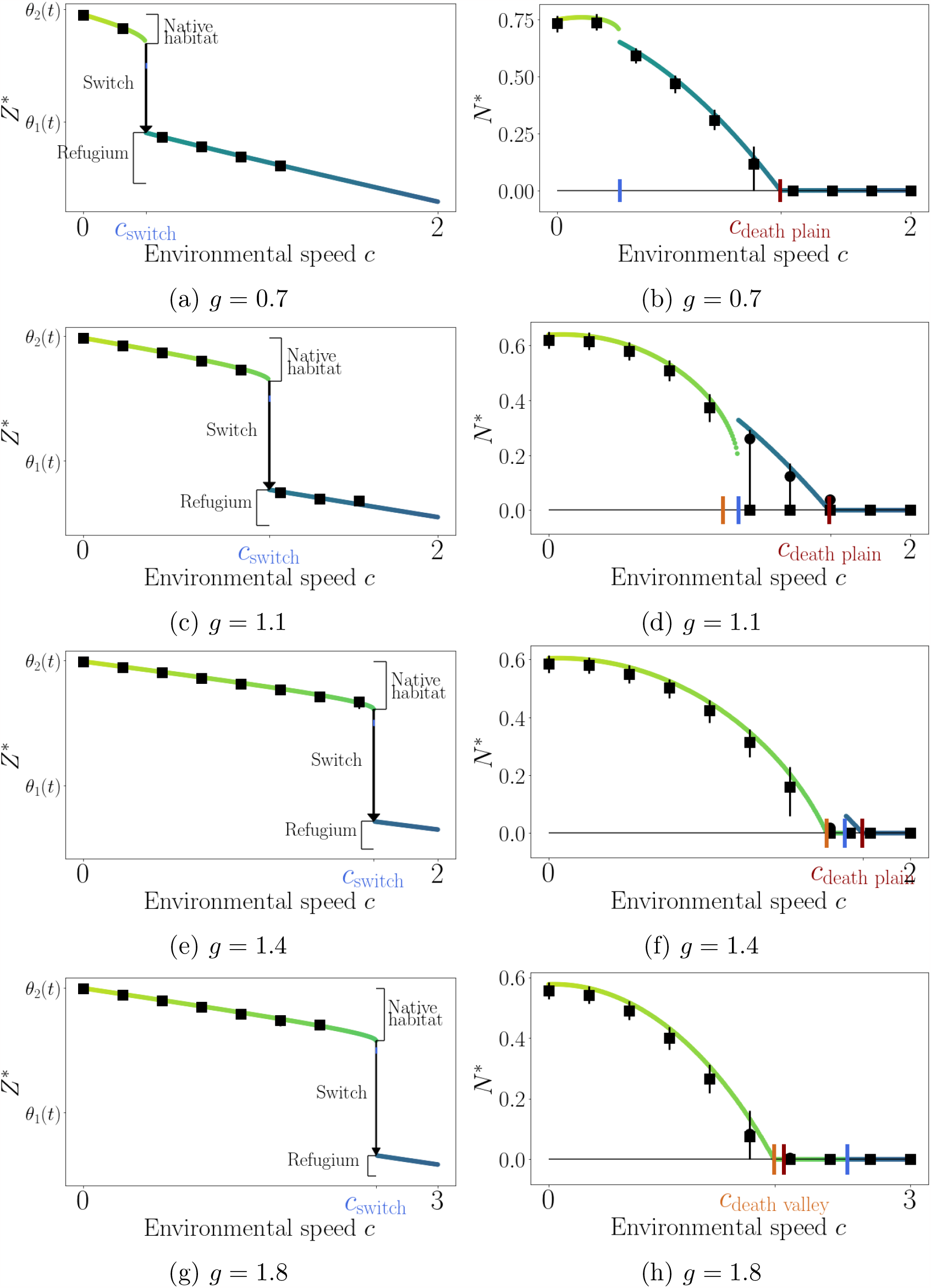
Same as Fig. 4 with final results of stochastic IBS instead of deterministic numerical resolutions. As in Fig. 4, the colored curves correspond to the analytical solution of Eq. (*S*) (green in the native habitat, blue in the refugium). The black squares represent the median quantities from the IBS and the vertical black lines span the results of 98% of the stochastic trajectories. Moreover, regarding the plots of *N* ^*^ (right column), the black circles indicate the median metapopulation size across non-extinct populations. Mean traits of extinct populations are ill-defined and thus not displayed.

**Figure 7.**
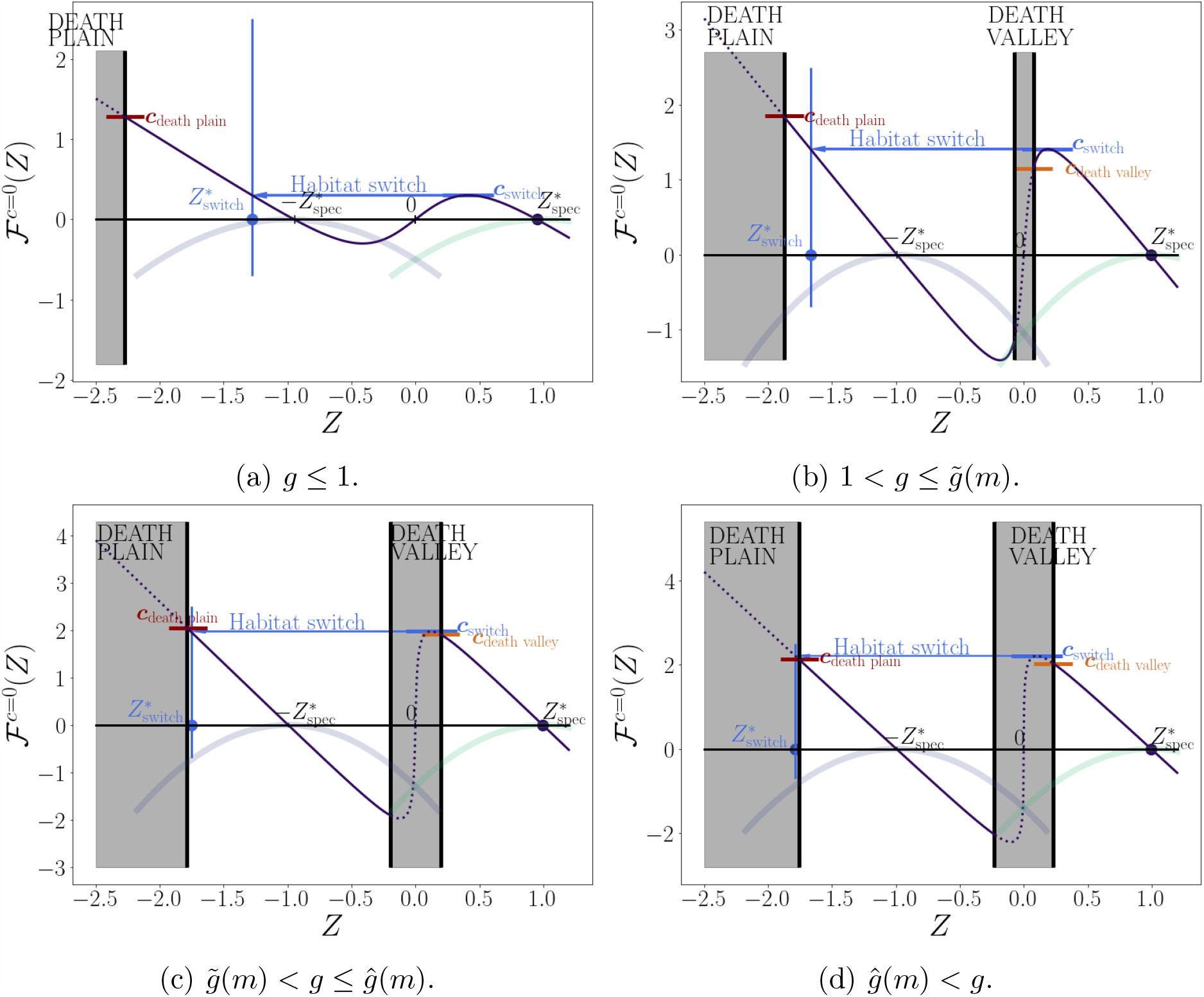
Same as Fig. 3, but with *m* = 0.2 and *g* = 0.5, 1.05, 1.3, 1.4 (top to bottom, left to right).

**Figure 8.**
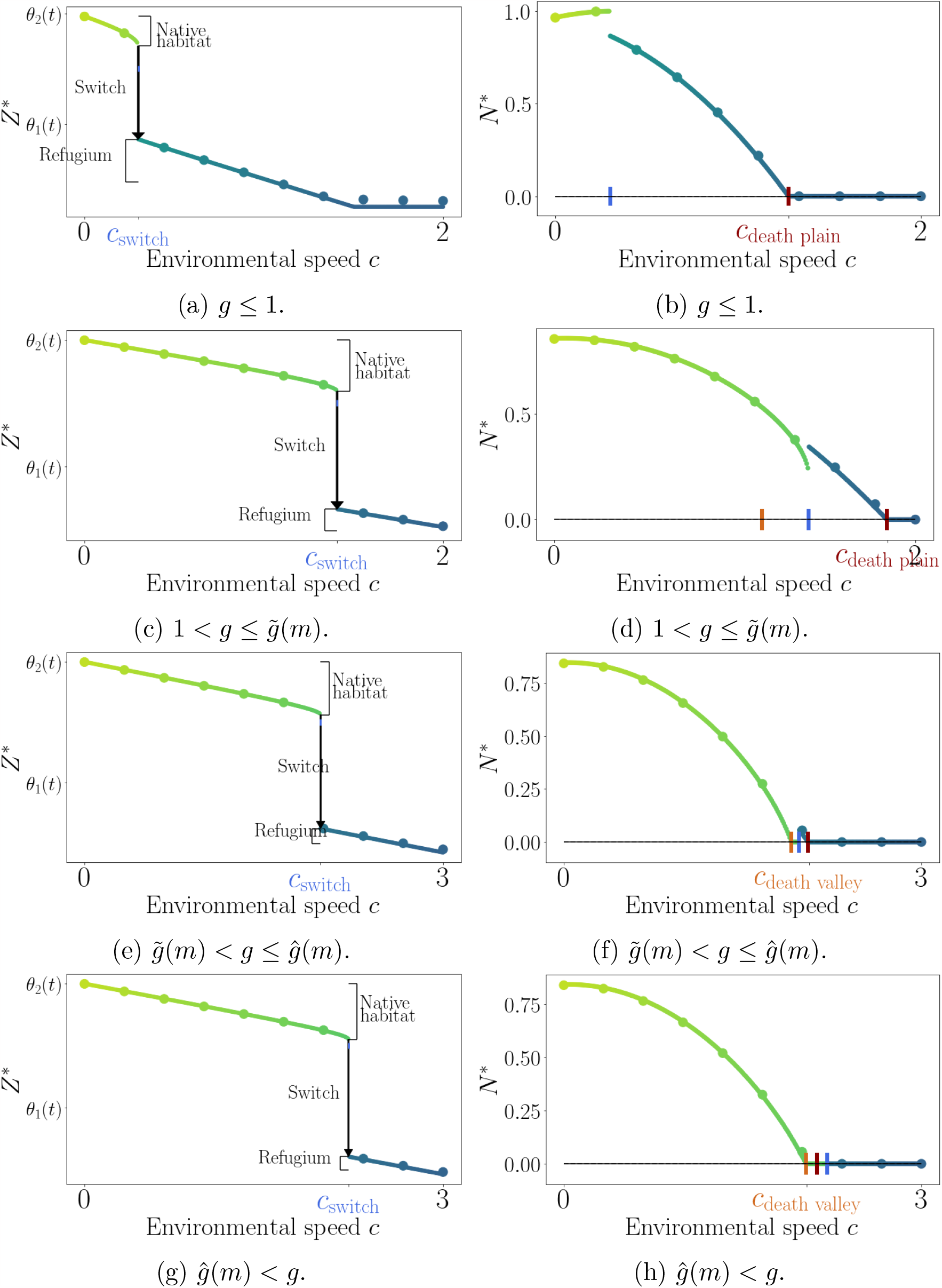
Same as Fig. 4, but with *m* = 0.2 and *g* = 0.5, 1.05, 1.3, 1.4 (top to bottom).

**Figure 9.**
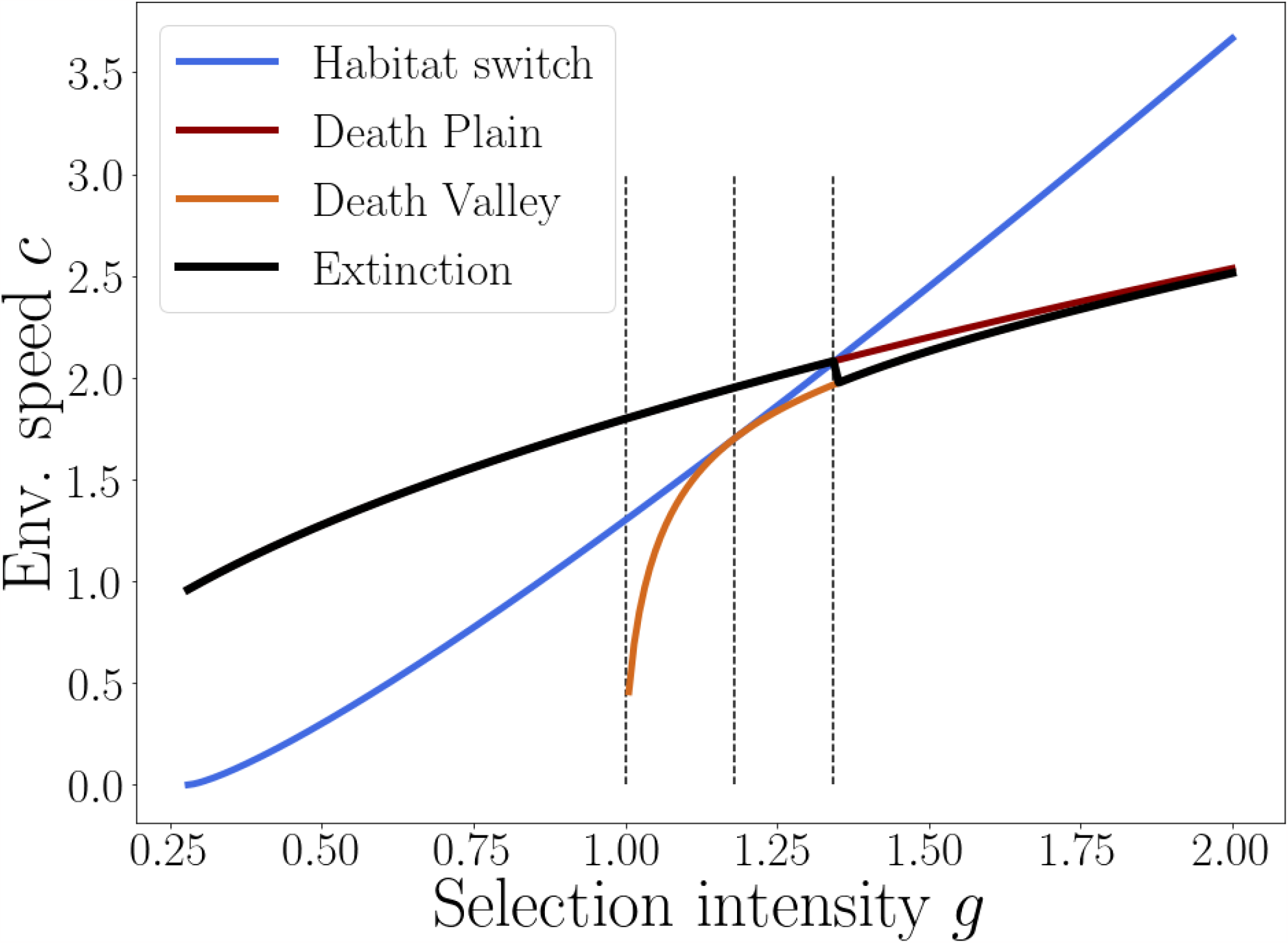
Same as Fig. 5, but with *m* = 0.2.

